# Auto-regulation of the real-time kinetics of the human mitochondrial replicative helicase

**DOI:** 10.1101/2024.11.14.623559

**Authors:** Ismael Plaza-G.A., María Ortiz-Rodríguez, Seth P. Buchanan, Samuel Míguez, Kateryna M. Lemishko, Fernando Moreno-Herrero, Rafael Fernandez-Leiro, Grzegorz L. Ciesielski, Borja Ibarra

## Abstract

The human mitochondrial replicative helicase, Twinkle, is essential for the replication and integrity of mitochondrial DNA (mtDNA). Therefore, investigating the real-time kinetics of Twinkle’s activities and their regulation is crucial for understanding mtDNA maintenance. Here, we combine biochemical and single-molecule manipulation and visualization techniques to investigate the loading of Twinkle onto the DNA fork, its real-time DNA unwinding and rewinding kinetics, and the regulation of these processes by the amino- (N) and carboxyl- (C) terminal ends of the helicase and the mitochondrial SSB protein (mtSSB). We observed that Twinkle rapidly diffuses along dsDNA scanning for the DNA fork, where it establishes specific interactions that halt further diffusion. Our results show that during DNA unwinding, the real-time kinetics of the helicase are downregulated by interactions of its N-terminal Zinc-binding domain (ZBD) with DNA and the control of the ATPase activity by the C-terminal tail. We found that binding of mtSSB to DNA likely outcompetes the ZBD-DNA interactions, alleviating the down regulatory effects of this domain. Furthermore, we show that ZBD-DNA interactions, together with the ATP binding, also control the real-time kinetics of the DNA rewinding events that follow helicase stalling. Our findings reveal that ZBD and C-terminal tail play a major role in regulation of Twinkle’s real-time kinetics. Their interplay constitutes an auto-regulatory mechanism that may be crucial for coordination of the mtDNA maintenance activities of the helicase.

## INTRODUCTION

Mitochondrial DNA (mtDNA) is maintained and replicated differently from its nuclear counterpart. Twinkle is the sole replicative helicase found in human mitochondria^1,2^. It presents strand separation activity needed for leading strand replication. In addition, Twinkle exhibits strand annealing, strand-exchange and branch migration activities and is essential for organization of mitochondrial RNA granules^3–7^, suggesting a multi-contextual role in mitochondrial nucleic acid metabolism. The essential function of Twinkle is underscored by disease variants of the helicase, which lead to alterations in the copy number and integrity of mtDNA associated with embryonic lethality and numerous heritable neuromuscular diseases^8–11^.

Twinkle belongs to the superfamily 4 (SF4) DNA helicases, which assemble into ring-shaped oligomers that bind NTPs at each of the subunits interfaces and DNA in the central channel^12–14^. Each subunit of Twinkle could be divided into an N-terminal domain (NTD) and a C-terminal domain (CTD), joined by a flexible helical linker (Figure 1A). The CTD of Twinkle, which shares high similarity with the CTD of the homologous bacteriophage T7 helicase^15^, contains the conserved helicase and ATPase motifs^16^.

**Figure 1:**
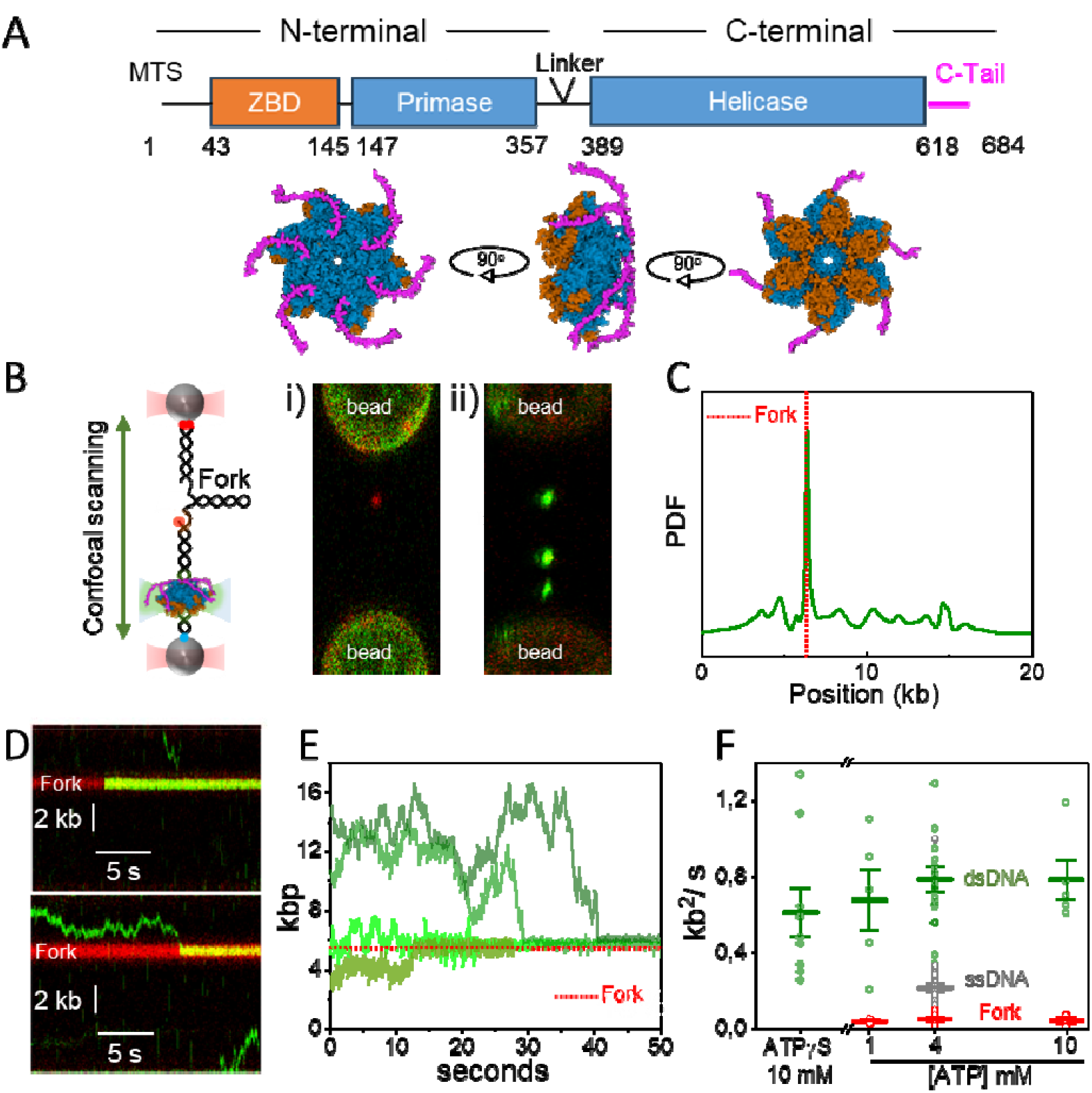
Loading and diffusion of Twinkle on DNA. **A)** (Top) Twinkle consists of an ancestral N-terminal primase domain and a C-terminal helicase domain, connected by a linker region. Residues (numbers) corresponding to the Zinc Binding Domain (ZBD, orange) and the carboxyl-terminal tail (C-Tail. Magenta) were deleted in the ΔZBD and ΔC-tail variants, respectively. MTS-Mitochondrial Targeting Sequence. (Bottom) Representation of Twinkle hexamer, with ZBD in orange and the C-tail in magenta, prepared by superimposing AlfaFold2^68^ predicted monomers with T7gp4 hexamer (PDB 1E0K). **B)** (Left) Diagram of the optical tweezers-confocal assay to image eGFP-Twinkle (green) on individual DNA constructs containing a DNA fork (446 bp, Fork) attached between optically trapped beads (Methods). (Right) Example scans showing separately i) the Atto 647-labelled fork position (red) and ii) Twinkle diffraction-limited spots (green) on the DNA construct. **C)** Distribution of initial positions of eGFP-Twinkle diffraction-limited spots on the DNA constructs containing a DNA fork (N= 50) showing preferential binding at the fork position. In contrast, eGFP-Twinkle did not present preferential binding positions on dsDNA molecules lacking a DNA fork (Figure S2). **D)** Representative kymographs of eGFP-Twinkle oligomers (green) binding directly (top) or diffusing before loading (bottom) at the fork position (in red). **E)** Representative position vs. time plots of eGFP-Twinkle spots on dsDNA with a forked structure show that, in all cases, diffusion is halted upon finding the fork position (4mM ATP). **F)** Diffusion coefficients of individual Twinkle units (6.0±2 monomers) on dsDNA (green, N=50), ssDNA (grey, N=38, Figure S4) and DNA fork (red, N=21) at different ATP concentrations or in the presence of ATPγS. Error bars represent standard error of the mean (s.e.).

These motifs are located between two adjacent CTDs, so that the binding of the NTPs effectively connects the subunits and enables their coordinated movement after NTP hydrolysis. The terminal residues of the CTD form an unstructured tail (C-tail) with no sequence similarity with the C-terminal tail of T7 helicase, but well conserved among most mammals^17^. Deletion of the C-tail increased NTP hydrolysis^12,17,18^, suggesting that the C-tail of the full-length helicase provides negative regulatory function. Twinkle NTD exhibits DNA binding activity, providing stability and uniformity to Twinkle oligomers^4,19,20^. Although the NTD contains primase motifs, this function has been lost in human Twinkle due to its deficiency to bind Mg^2+^ in the ancestral catalytic site^15^. The terminal zinc-binding domain (ZBD) confers ssDNA binding activity, which in the prototypical T7 helicase contributes to the priming process^21^. In humans, however, ZBD appears to lack three of the four conserved cysteine residues that coordinate Zn^2+^ in the T7 helicase^15,22^, and its function remains unknown.

Structural and biochemical data showed that Twinkle can exist in various oligomeric states in solution, from broken rings to higher order ≥6-mer ring-like oligomers^12,13,22,23^. Twinkle binds both single-stranded and double-stranded (ds)DNA^8,24,25^ and can self-load onto single-stranded (ss)DNA circles^8,20,25^. Recent AFM and HS-AFM studies on human and *Lates calcarifer* Twinkle (LcTwinkle) have shown that Twinkle ring-like oligomers can switch between close-to-open conformations, allowing them to load (and unload) onto dsDNA without the need for a dedicated helicase loader^26,27^. The dual ability of Twinkle to load onto both ds- and ss-DNA raises the question of how the helicase recognizes the DNA fork position within the ∼16 kbp long mtDNA.

Upon loading at the DNA fork, Twinkle catalyzes the unwinding of duplex DNA in a 5’ to 3’ direction, driven by the hydrolysis of nucleoside triphosphate^4,24,28^. Recent biochemical and single-molecule studies suggest the existence of multiple ssDNA-binding sites on the helicase surface^3,5,19^, supporting a steric exclusion and wrapping (SEW) model for DNA unwinding^29^. Interestingly, Twinkle exhibits a DNA unwinding activity limited to a few base pairs (20-40 bp)^4,24,28^, which is enhanced to kilo base pairs processivity by its partners at the replisome^24,28^; Polγ and the mitochondrial SSB protein (mtSSB). This behavior suggests a strong autoregulation. However, the origin of autoregulation and the mechanism by which is relieved by other replication components remain unexplored.

In this study, we combined biochemical and single-molecule manipulation and visualization techniques to interrogate the effects of ATP, helicase-DNA interactions and mtSSB on the loading and real-time DNA unwinding and rewinding kinetics of the human mitochondrial helicase Twinkle. We compared wild-type helicase (WT) activities with those of truncation variants at the N- (ΔZBD) and C-terminal (ΔC-tail) ends to assess their putative regulatory roles on the helicase activities. The ΔZBD variant lacks the first 145 residues of the N-terminal domain, which correspond to the ZBD in the homologous T7 helicase^2,21^, Figure 1A. These residues stabilize binding to the translocation strand without affecting the ATPase activity of the helicase significantly^20^. The ΔC-tail variant lacks 66 residues of the unstructured C-terminal tail, Figure 1A. This truncation increases ATPase activity without altering DNA binding significantly^9,12,17,18^.

Our results show that Twinkle can rapidly diffuse on long dsDNA scanning for the DNA fork. Additionally, interactions between Twinkle’s C-tail and mtSSB assist greatly in the functional loading (i.e. loading followed by unwinding) of the helicase at the fork. During DNA unwinding, the real-time kinetics of the helicase are efficiently auto-regulated by ZBD-DNA interactions and the control of ATPase activity by the C-terminal domain. Binding of mtSSB to ssDNA competes for ZBD-DNA interactions, alleviating the suppressive effects of this domain. Furthermore, our results indicate that the ZBD and C-tail also control the DNA rewinding events characteristic of Twinkle activity following DNA unwinding.

## RESULTS

### Rapid diffusion of Twinkle on dsDNA facilitates fork recognition

First, we investigated how Twinkle recognizes a single DNA fork within a long dsDNA compatible with the length of the mtDNA (∼16 kbp). We used optical tweezers combined with confocal scanning microscopy and microfluidics^30^ to image the position of fluorescent Twinkle (eGFP-Twinkle) oligomers in real-time along individual 18 kbp-long dsDNA molecules containing a single DNA fork (446 bp), tethered under constant tension (3 pN) between two optically trapped beads (Figure 1B, left panel, Methods). The fork was labelled with Atto647 at the 3’-end to locate its position (Figure 1B, panel i) and contains a 32 nt long 5’-ssDNA gap to facilitate Twinkle DNA unwinding activity (Methods). Upon DNA attachment, eGFP-Twinkle (1 nM) was flown into the reaction chamber and detected as diffraction-limited spots on the DNA (Figure 1B panel ii). Of note, eGFP-Twinkle unwinds DNA with processivity and average velocity identical to those of the wild-type (WT) helicase at the single-molecule level (Figure S1).

In the absence of ATP, a small fraction (∼2.5%) of the fork DNA constructs contained diffraction-limited fluorescent spots corresponding to eGFP-Twinkle. In sharp contrast, in the presence of 4 mM ATP, all trapped DNA molecules contained one or several diffraction-limited fluorescent spots (Figure 1B panel ii) indicating that ATP favored loading of the helicase on individual dsDNA molecules without free-ends. We observed a wide distribution of initial binding positions with a predominant peak at the location of the DNA fork, which represents just ∼0.16% of the total length of the trapped DNA construct (Figure 1C). Fluorescence spots bound to the DNA fork remained apparently static (Figure 1D). However, those bound to the dsDNA moved bidirectionally upon finding the protein-free DNA fork, where movement was halted (Figures 1D and 1E). We checked that on dsDNA molecules lacking the DNA fork, eGFP-Twinkle does not present preferential binding and its diffusion is not halted at specific locations (Figure S2). These results show that loading of the helicase at the fork can be attained alternatively after fast scanning of the dsDNA.

The number of Twinkle monomers per diffraction-limited spot was estimated by normalizing their fluorescent intensity by that of a control eGFP-protein fusion (Methods and Figure S3). We considered oligomeric assemblies containing 6.0±2 eGFPs as individual Twinkle units, whereas multiples of this value were considered either partial (broken rings) or higher oligomers (Figure S3). Analysis of the motion of diffraction-limited spots compatible with one unit of Twinkle revealed that they presented an average diffusion coefficient on dsDNA (*D*_*ds*_) of 0.075 ± 0.006 *μ*m^2^/ s (N=20), which corresponds to 0.787 ± 0.065 kb^2^/ s (Figures 1F and S3, Methods). Almost identical diffusion coefficients were measured at different ATP concentrations and in the presence of the non-hydrolysable ATP analog ATPγS (N=30), showing that diffusive movement did not depend on ATP hydrolysis, Figure 1F. Because the inner diameter of Twinkle oligomers (>6-mer) is wide enough to accommodate dsDNA in the central channel^13,14^, we presume that the rapidly diffusive Twinkle is topologically linked to dsDNA. In fact, the measured *D*_*ds*_ values are in line with those reported for the diffusion coefficients of other eukaryotic proteins known to encircle dsDNA, such as the Fanconi anemia D2-I complex^31^ and CMG helicase^32,33^. In sharp contrast, Twinkle remained static at the DNA fork position at all ATP concentrations during the observation time, or eGFP-bleaching time (∼120 seconds). The average diffusion coefficient at the fork (*D*_*fork*_) was ∼0.04 kb^2^/ s (N=21) at all ATP concentrations tested (Figure 1F), which probably provides a lower limit on the measurable diffusion constant under our imaging conditions. In an independent set of control experiments, we measured that Twinkle units also diffuse on ssDNA with a diffusion coefficient (*D*_*ss*_) of 0.21 ± 0.02 kb^2^/ s (N=38, 4 mM ATP), Figures 1E and S4. This rate is 5-times faster than that of Twinkle on the DNA fork, indicating that the static binding at the forked position is not caused by interaction of Twinkle with the ssDNA portion of the fork. Instead, interactions with the two strands of the DNA fork are more likely required to stall diffusion of the helicase.

### Modulation of Twinkle’s real-time DNA unwinding kinetics by ATP binding

We used dual-beam counter-propagating optical tweezers^34^ to monitor the real-time kinetics of wild-type and two deletion variants of Twinkle on individual forked-like DNA constructs (Methods), Figure 2A. DNA unwinding traces were measured at constant tension from the increase in the end-to-end extension of the tether as each base pair of the DNA hairpin is converted to two single-stranded nucleotides (Methods and Figures 2A and 2B). In many cases, unwinding activities were followed by DNA rewinding events detected as a reduction in the end-to-end length of the tether, Figure 2B. We analyzed unwinding and rewinding events independently, and we will discuss them here sequentially.

**Figure 2:**
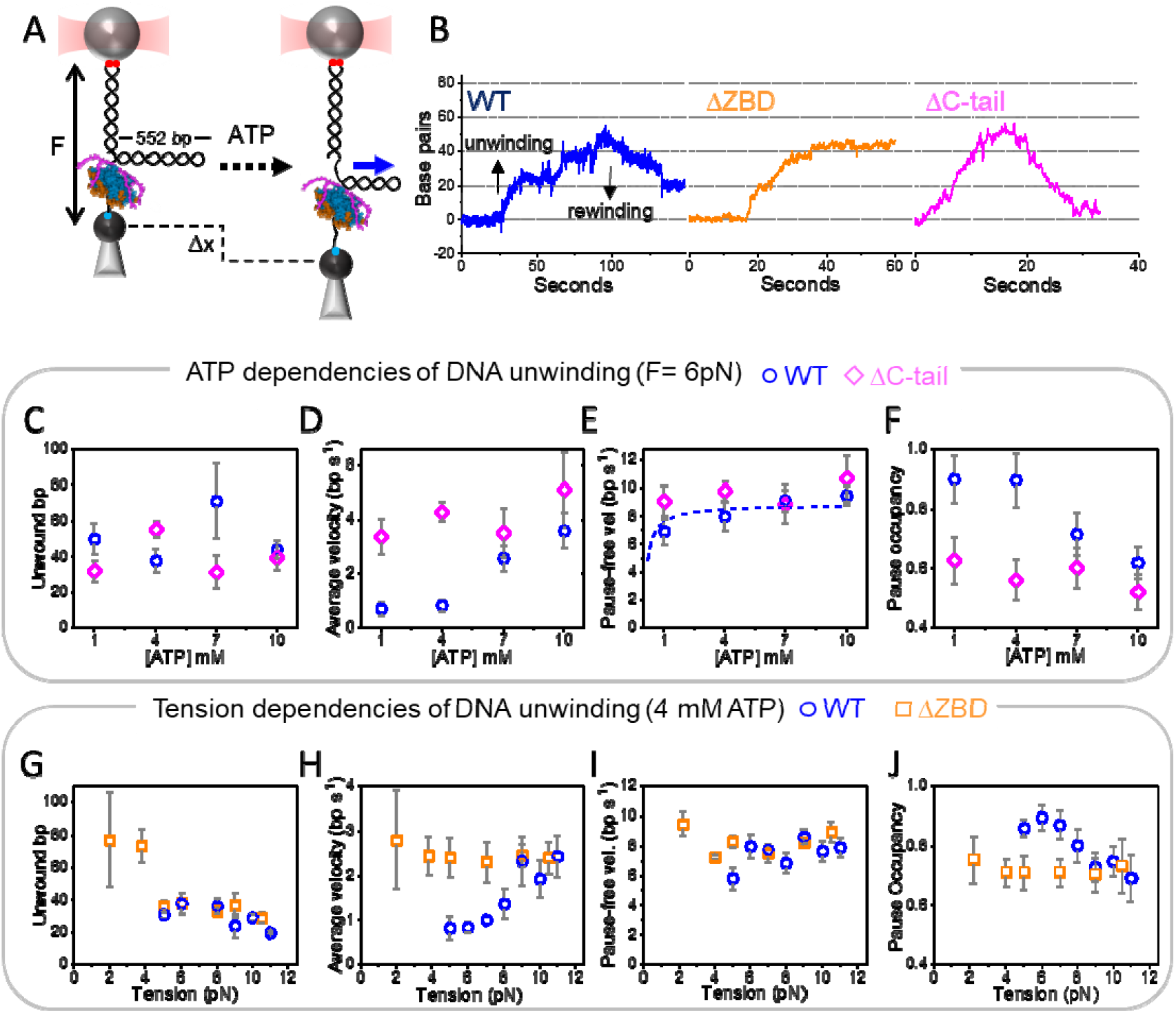
ATP and tension dependencies of the real-time DNA unwinding kinetics. **A)** Schematic of optical trapping assays. A DNA fork-like construct is tethered between two functionalized micron-sized beads. The construct consists of a DNA hairpin (552 bp) connected to a bead in the optical trap (red cone) via a ∼2.6-kb long dsDNA handle and to a bead on a micropipette through a poly-(dT)_35_ that serves as a helicase loading site (Methods). In the presence of ATP, DNA unwinding was recorded as a change in tether extension (Δx) at constant mechanical tension (F). Blue arrow indicates 5’ to 3’ translocation of the helicase along the hairpin. **B)** Representative traces of the WT Twinkle (blue), ΔZBD (orange) and ΔC-tail (magenta) variants (F=6 pN). In many cases (see main text), DNA unwinding (bp increase) was followed by rewinding (bp decrease). **C** to **F)** ATP dependencies of the average unwinding processivity (**C**), average unwinding rate (**D**), pause-free velocity (**E**) and pause occupancy (**F**) for the WT helicase (blue), and ΔC-tail (magenta) variants. Dotted line in E shows the sigmoidal increase of the WT Twinkle pause-free velocity with ATP concentration (see Figure S5 for additional data and fits). For plots C-F data was recorded at F=6 pN, the minimum tension that consistently allowed detection of activities at all ATP concentrations. **G** to **J)** Tension dependencies of the average unwinding processivity (**G**), average unwinding rate (**H**), pause-free velocity (**I**) and pause occupancy (**J**) for the WT helicase (blue) and the ΔZBD variant (orange). Error bars in all data plots represent s.e.. Table 1 summarizes the ATP and tension dependencies of the unwinding kinetics of the three helicases under study.

Similar to other helicases^35–39^, ATP turnover is expected to regulate the real-time kinetics of DNA unwinding by Twinkle. Initially, we examined the effect of varying ATP concentrations (from 1 mM to 10 mM) on the real-time kinetics of DNA unwinding of wild-type (WT) Twinkle (5 nM, tension F=6 pN, number of molecules N=55, Figures 2C-F). Analysis of individual DNA unwinding traces revealed that the average unwinding processivity did not change with ATP concentration significantly (Figure 2C, Table 1). The measured unwinding processivity, ∼40 bp, was more than ten-times shorter than the length of the DNA fork (552 bp) and is consistent with previous biochemical studies of this helicase^4,24,28^. In contrast, the increase of ATP concentration from 1 to 10 mM stimulated the average DNA unwinding rate 5-fold, from ∼0.8 bp/ s to ∼4 bp/ s, Figure 2D and Table 1. The average unwinding rate averages out active unwinding burst with pauses, or transient inactive states, which frequently interrupted the helicase advancement (Figure 1B). To discern the effect of ATP concentration on the active and pause states, we calculated the pause-free unwinding rate^40^ and the pause occupancy (Methods). The latter was defined as the ratio between the average rate and the pause-free rate and estimates the probability of finding the helicase in a pause state. This analysis showed that while the pause-free rate presented the expected sigmoidal increase (Figures 2E and S5), the pause occupancy decreased almost 2-fold with increasing ATP concentrations (Figure 2F and Table 1). These findings point to ATP binding as a strong regulator of the characteristic high pause occupancy of Twinkle, which in turns would modulate the unwinding kinetics of the helicase at near physiological ATP concentrations^41,42^.

**Table 1.** Tension and ATP concentration dependencies of average unwound nucleotides, average unwinding velocities (bp/s) and pause occupancies of the helicases under study. Tension dependencies were assayed at 4 mM ATP both, in the absence and presence of mtSSB. Upper indexes indicate the minimum tension (Low) at which values were recorded. The maximum tension (High) was in all cases 11 or 12 pN. ATP concentration dependencies (from 1 to 10 mM) were recorded at 6 pN of tension. Arrows indicate the change in the values as a function of mechanical tension or ATP concentration. Numbers correspond to rounded values.

|  |  | Tension<br>Low→High |  | ATP<br>1→10 mM |
| --- | --- | --- | --- | --- |
|  |  | No mtSSB | With mtSSB | No mtSSB |
| Average<br>unwound bp | WT | 40<sup>6pN</sup> →20 | 130<sup>2pN</sup> →30 | 40 →40 |
|  | ΔZBD | 80<sup>2pN</sup> →20 | 80<sup>2pN</sup> →20 | 40 →40 |
|  | ΔC-tail | 45<sup>6pN</sup> →30 | 45<sup>6pN</sup> →30 | 40 →40 |
| Average<br>velocity(bp/s) | WT | 0.8<sup>6pN</sup> →2.5 | 2.5<sup>2pN</sup> →0.8 | 0.8 →4 |
|  | ΔZBD | 2.5<sup>2pN</sup> →2.5 | 2.5<sup>2pN</sup> →2.5 | 1.5 →4 |
|  | ΔC-tail | 4<sup>6pN</sup> →4 | 4<sup>6pN</sup> →4 | 4 →5 |
| Pause<br>occupancy | WT | 0.9<sup>6pN</sup> →0.7 | 0.7<sup>2pN</sup> →0.9 | 0.9 →0.6 |
|  | ΔZBD | 0.7<sup>2pN</sup> →0.7 | 0.7<sup>2pN</sup> →0.7 | 0.8 →0.5 |
|  | ΔC-tail | 0.6<sup>6pN</sup> →0.6 | 0.6<sup>6pN</sup> →0.6 | 0.6 →0.5 |

In line with these findings, we observed that the Twinkle C-terminal tail deletion variant, which exhibits enhanced ATPase activity *in vitro* (ΔC-tail, 5 nM, N=36), showed similar average processivity but a 5-times faster average unwinding rate than the WT helicase at ≤4 mM ATP (Figures 2C and 2D and Table 1). The deletion of the C-tail did not affect significantly the pause-free velocity of the variant with respect to that of the WT (Figure 2E), showing that its increased average unwinding rate is mainly due to a reduced pause occupancy (Figure 2F and Table 1). At ATP concentrations ≤4 mM, pause occupancy of the ΔC-tail variant was about half that of the WT. Accordingly, we showed that this variant is 2-5-fold more effective than the WT helicase in DNA unwinding *in vitro* (Figures S6A and S6B). Overall, these findings suggest that the C-terminal tail hinders ATP turnover, which in turn, would increase pause occupancy slowing down the DNA unwinding kinetics of Twinkle.

### Modulation of the real-time DNA unwinding kinetics by helicase-fork interactions

Helicase-DNA interactions are crucial to couple ATP turnover with DNA unwinding and mechanical tension applied to the opposite strands of the DNA fork affect these interactions^43–46^. Thus, we aimed to determine the role of Twinkle-fork interactions on DNA unwinding by measuring the response of the helicase DNA unwinding kinetics to increasing mechanical tension (Figure 2A). Measurements were carried out over a range of constant tension below that required to unfold the hairpin mechanically (∼12 pN, Methods) at near physiological ATP concentration (4 mM ATP).

Individual DNA unwinding traces of WT Twinkle were detected at a minimum tension of 5 pN, with an average processivity of ∼40 bp (N= 47, Figure 2G and Table 1). Interestingly, processivity was found to decrease slightly with increasing tensions, Figure 2G. In contrast, tension stimulated by ∼3-fold the average unwinding rate from ∼0.8 bp/ s to ∼2.5 bp/ s (Figure 2H, Table 1). Because, tension had no significant effect on the pause-free rate (∼8 bp/ s, Figure 2I), stimulation of the average velocity could be attributed to the effect of tension on decreasing the pause occupancy of the helicase, from ∼90% (5pN) to ∼70% (11 pN) (Figure 2J, Table 1). This stimulatory effect of mechanical tension could be interpreted as if tension would disrupt helicase-fork interactions that slow down the average velocity. However, we cannot discard the possibility of tension favoring the ATPase activity of the helicase.

To distinguish between these two possibilities, we measured the activity of the Twinkle deletion variant ΔZBD, which exhibits similar ATPase activity but reduced DNA binding compared to the WT Twinkle^20,23^. Initially, we corroborated that under our experimental conditions the DNA unwinding kinetics of ΔZBD depend on ATP concentration in a manner similar to that of the WT helicase, as expected for this variant (Figure S7, N=46). Next, we measured the effect of mechanical tension on the real-time kinetics of the variant. Individual DNA unwinding traces of ΔZBD were detected consistently at tension as low as 2 pN (3 pN lower than that required to detect WT activity) and with an enhanced average processivity of ∼80 bp (Figure 2G). These results indicate that elimination of ZBD favors functional loading (loading followed by initiation of DNA unwinding) at the fork and promotes processivity. As in the case of the WT, the average processivity of the ΔZBD variant decreased gradually with tension (Figure 2G). In addition, the average unwinding rate of ΔZBD was 2-3 times faster, and its pause occupancy lower, than those of the WT Twinkle (< 8 pN) and they did not show any significant tension dependencies (Figures 2H and 2J). These results go in line with *in vitro* biochemical assays showing an enhanced DNA unwinding activity of the ΔZBD variant (Figure S6) and overall, point to the ZBD as a source of pauses during DNA unwinding. The fact that tension did not promote the average rate of the ΔZBD variant up to values found at higher ATP concentrations, suggested that tension does not have an effect on the ATPase activity of the enzyme. Therefore, the observed dependency of the WT helicase average unwinding rate with tension could be attributed to the (known) effect of tension on affecting helicase-fork interactions. A putative scenario compatible with our data is that tension would disrupt the interactions established by ZBD of the WT helicase and the DNA, which otherwise, would downregulate DNA unwinding rate by favoring pause occupancy at near physiological ATP concentrations.

### mtSSB binding mode stimulates the real-time DNA unwinding kinetics of WT Twinkle

mtSSB is known to stimulate the DNA unwinding activity of Twinkle *in vitro*^18,24,26^. However, the specific mechanism behind this stimulation remains unclear. To investigate this, we measured the effect of mtSSB (5 nM) on the tension-dependent real-time kinetics of WT Twinkle and the two variants used in this study (at 4 mM ATP, Figure 3A and 3B, Table 1). In the case of the WT, mtSSB favored the detection of DNA unwinding activities at tension 3 pN lower than that in the absence of mtSSB. This result indicates that mtSSB binding to the 5’ ssDNA tail favors the functional loading of the helicase at the fork. This in agreement with the stimulation of Twinkle DNA unwinding activity by mtSSB, as measured in our bulk biochemical assays, Figure S6C. At the lowest tension (2 pN), mtSSB stimulated by ∼3-4 times the average processivity and DNA unwinding rate of the WT helicase with respect to the values expected at this tension in the absence of mtSSB, Figures 3C and 3D. These results argue that the increase in processivity is a direct consequence of the increase in the average DNA unwinding rate. The stimulus in the average velocity aligns well with the effects of mtSSB and RPA on Twinkle and CMG DNA unwinding rates measured previously *in vitro*, respectively^26,33^. Our results showed that the stimulation of average velocity was due to the effect of mtSSB on decreasing the pause occupancy rather than to increasing the pause-free velocity, which remained unaltered (Figures 3E and 3F). These stimulatory effects on the real-time kinetics of DNA unwinding seemed specific to mtSSB because significant stimulation was not detected with identical concentrations of the homologous *E. coli* SSB (Figure S8). Interestingly, the stimulatory effect of mtSSB on the helicase kinetics ceased as tension increased; stimulation was no longer detected above ∼6 pN (Figures 3C, 3D and 3F). Tension reduces the binding mode or binding footprint of mtSSB to ssDNA^47,48^, suggesting that the SSB binding mode is relevant for stimulation of helicase activity.

**Figure 3:**
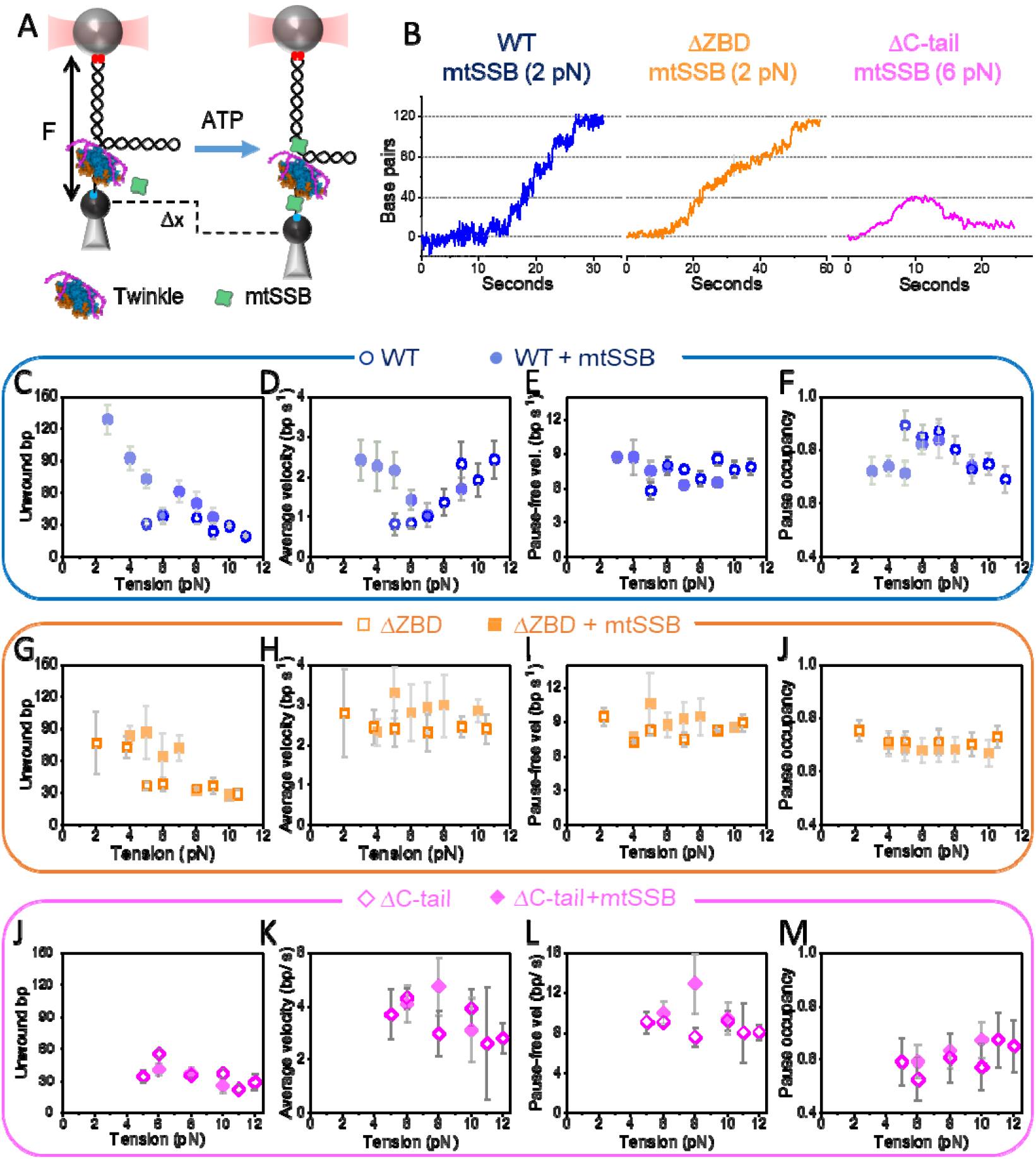
Tension dependencies of the real-time DNA unwinding kinetics in the presence of mtSSB. **A**) Schematic of optical trapping assays in the presence of the helicase and mtSSB. Experimental design is identical to that described in Figure 2A with the exception of including mtSSB (5 nM, green shape) in the reaction buffer. Experiments were carried out at 4 mM ATP. **B)** Representative data traces in the presence of mtSSB for the WT Twinkle (blue, F= 2 pN), ΔZBD (orange, F=2 pN) and ΔC-tail (magenta, F= 6 pN) variants. Under these conditions, only the ΔC-tail activities showed significant rewinding events upon unwinding. **C-F)** Tension dependencies of the average unwinding processivity (**C**), average unwinding rate (**D**), pause-free velocity (**E**) and pause occupancy (**F**) for the WT helicase in the absence (blue empty symbols) and presence (solid blue symbols) of mtSSB. Shaded green area highlights the tension range at which mtSSB stimulatory effects are apparent. **G-J)** same as above (C to F) but for the ΔZBD variant in the absence (empty orange symbols) and presence (full orange symbols) of mtSSB. **J-M)** same as above (C to F) but for the ΔC-tail variant in the absence (empty magenta symbols) and presence (solid magenta symbols) of mtSSB. Error bars in all data plots represent s.e.

In contrast, the binding of mtSSB to DNA did not favor the detection of activities at lower tension, nor did it have a significant impact on the DNA unwinding kinetics of the ΔZBD and ΔC-tail Twinkle variants at any mechanical tension, compared to conditions when it was absent (Figures 3G-M). Biochemical analysis further confirmed that mtSSB does not stimulate the DNA unwinding activity of Twinkle variants (Figure S6C). These results highlight the relevance of the ZBD and C-tail on mediating functional and/or physical interactions with the mtSSB that would facilitate functional loading of the helicase onto the DNA fork and/or promote DNA unwinding kinetics.

### ATP binding and ZBD-DNA interactions control rewinding events

Single-molecule experiments revealed that many individual unwinding traces were followed by rewinding events (Figure 2B). We did not observe in any case rapid or instantaneous rewinding of the DNA fork itself, which would suggest fast helicase slippage or detachment. Interestingly, significant differences were observed in the rewinding probabilities and kinetics of the WT and helicase variants under study, depending on ATP concentration and mechanical tension (Figures 4A-F).

**Figure 4:**
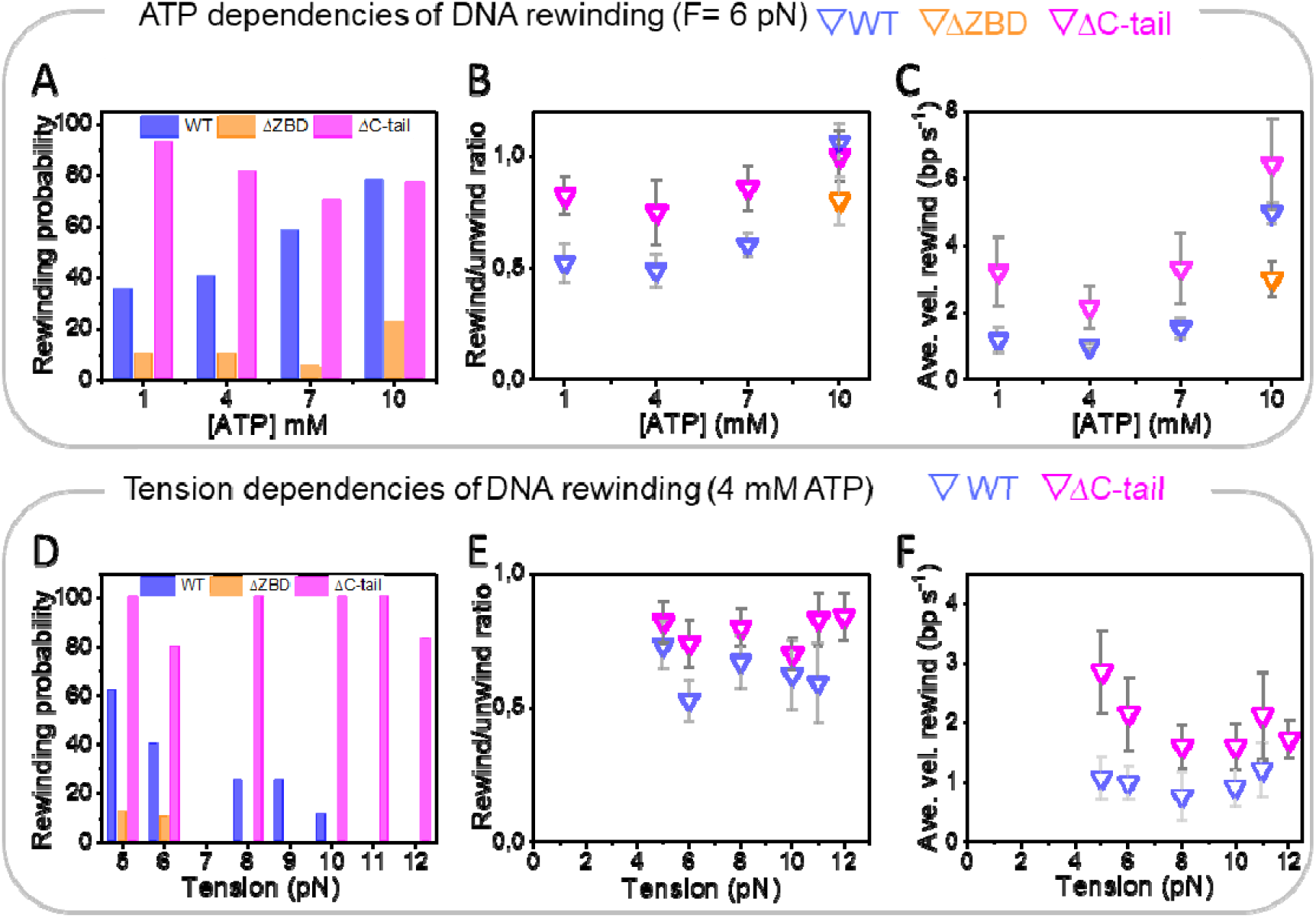
ATP and tension dependencies of DNA rewinding events. **A)** Probabilities of occurrence of rewinding events following unwinding stalling as a function of ATP concentrations for the WT helicase (blue bars), ΔZBD (orange bars) and ΔC-tail (magenta bars) variants. **B)** Average ratios between the total number of rewound and unwound bp as a function of ATP concentration for the WT (blue symbols), ΔZBD (orange symbol) and ΔC-tail (magenta symbols) variant. Ratios close to 1 indicate that the number of rewound bp is similar to the number unwound bp. **C)** Average rewinding velocities of the WT (blue symbols), ΔZBD (orange symbol) and ΔC-tail (magenta symbols) variants. B) and C) show the data for the ΔZBD variant only at 10 mM ATP because, below this concentration, the low number of rewinding events hampered the significance of the data analysis. **D)** Probabilities of occurrence of rewinding events following unwinding stalling as a function of tension for the WT helicase (blue bars), ΔZBD (orange bars) and ΔC-tail (magenta bars) variants. **E)** Average rewinding/ unwinding ratios as a function of tension for the WT (blue symbols), and ΔC-tail (magenta symbols) variant. **F)** Average rewinding velocities of the WT (blue symbols) and ΔC-tail (magenta symbols) variant. ATP and tension dependencies were measured at 6 pN and 4 mM ATP, respectively. Error bars in all data plots represent s.e.

In the case of the WT helicase, the probability of occurring of rewinding events increased from ∼35 to ∼80% with ATP concentration (Figure 4A, F=6 pN). These rewinding probabilities are in good agreement with those reported previously for WT Twinkle at comparable ATP concentrations by single-molecule FRET studies^5^. The number of rewound base pairs was generally smaller than that of unwinding, *i*.*e*. for WT Twinkle the average rewinding/unwinding ratio (i.e. average number of rewound base pairs divided by the number of unwound base pairs) was ∼0.5 at <10 mM ATP, Figure 4B. This ratio, along with the average velocities of rewinding, exhibited increasing trends with ATP concentration (Figures 4B and 4C). These ATP dependencies indicate that ATP turnover favors the rewinding events. In line with this hypothesis is that the ΔC-tail variant (with augmented ATPase activity) presented higher rewinding probabilities (∼80-100%), rewinding/unwinding ratios and rewinding rates than those of the WT at ATP concentrations below 10 mM, Figures 4A-C. In sharp contrast, the rewinding probability of the ΔZBD variant was just ∼5-10% at <10 mM ATP and only increased to 20% at 10 mM (Figure 4A). These results indicate that the ZBD domain and/or ZBD interactions with the DNA are essential to promote rewinding events.

Next, we showed that mechanical tension above 5-6 pN had no significant effect on the average unwinding/rewinding ratios and rates, but it did affect the rewinding probabilities of WT Twinkle and ΔC-tail variant differently (Figures 4D-F). In the case of the WT, mechanical tension decreased the occurrence of rewinding events from 60% (5 pN) to 0% (11-12 pN), Figure 4D. These results suggest that mechanical tension would disrupt WT helicase-DNA interactions relevant for rewinding. In line with this hypothesis, we measured that the presence of mtSSB (and EcoSSB) inhibited the occurrence of the WT rewinding events at all tensions, suggesting mtSSB would outcompete the WT helicase for DNA interactions relevant for rewinding. In sharp contrast, mechanical tension did not affect the rewinding probability of the ΔC-tail variant, Figure 4D. This result highlights again that elevated ATP turnover would favor rewinding events even if helicase-DNA interactions are compromised, as in this case, by application of mechanical tension to the DNA fork. Additional evidence further support this observation: *i*) the rewinding probability of the WT did not depend on tension when ATP concentration was increased to 10 mM (Figure S9). *ii*) mtSSB did not inhibit the rewinding probability of the ΔC-tail variant at any tension (Figure S9). Altogether, our data indicates that helicase-DNA interactions, probably mediated by ZBD, together with ATP binding, control rewinding events.

## DISCUSSION

Here, we presented single-molecule studies supported by biochemical assays to investigate Twinkle’s DNA fork recognition and unwinding kinetics. Table 1 summarizes the measured ATP and tension dependencies and Figure 5 outlines our main results. Briefly, we showed that Twinkle rapidly diffuses on long dsDNA molecules and scans for a DNA fork where it binds stably. Additionally, functional loading at the fork is favored by interactions between Twinkle’s C-tail and mtSSB. We also found that Twinkle is a strongly auto-regulated helicase. Its real-time kinetics of DNA unwinding are slowed down by high pause occupancy, which is caused by ZBD-DNA interactions and the control of ATPase activity by the C-terminal domain. Binding of mtSSB to ssDNA competes for ZBD-DNA interactions, alleviating the restraining effects of this domain. Furthermore, our results indicate that the ZBD and C-tail also control the DNA rewinding events characteristic of Twinkle activity upon DNA unwinding.

**Figure 5:**
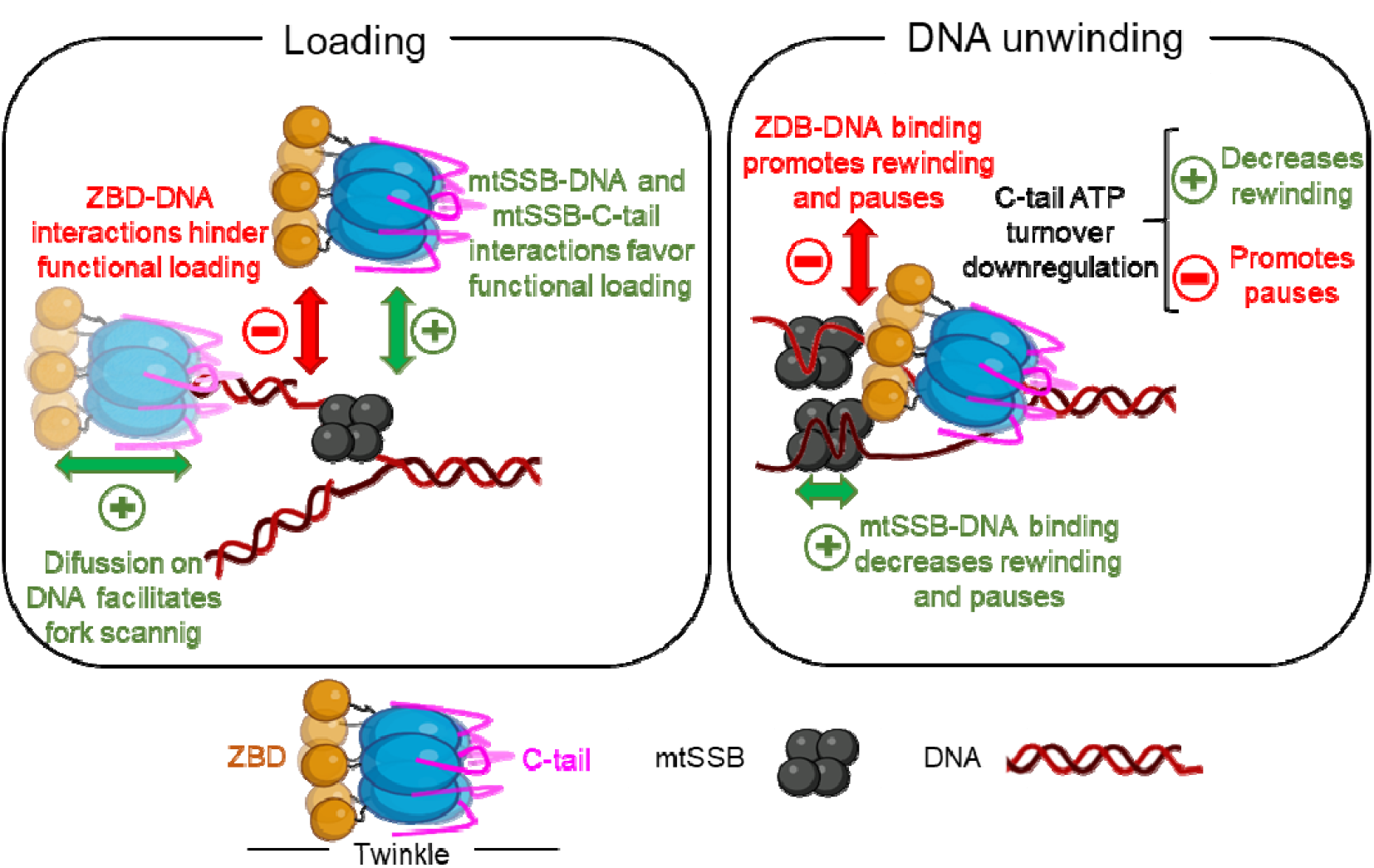
Auto-regulation of Twinkle activities by the ZBD and C-terminal tail. Diffusion along the DNA facilitates the rapid scanning of Twinkle for the fork junction. In the absence of mtSSB, ZBD-DNA interactions restrain functional loading at the fork and promote pauses (and rewinding events) during DNA unwinding. During this reaction, the C-tail downregulates ATP turnover, promoting additional pauses and decreasing DNA unwinding rate. In the presence of mtSSB, interactions between the mtSSB and the C-terminal tail of the helicase favor its functional loading onto the fork. Furthermore, mtSSB binding to the translocation strand may outcompete the suppressive ZBD-DNA interactions, favoring functional loading, decreasing pause occupancy during DNA unwinding, and reducing the probability of rewinding.

Loading of replicative helicases onto the DNA fork is often required for replisome assembly and therefore, a critical regulatory step in DNA replication^49,50^. Our data supports that Twinkle can diffuse on dsDNA for thousands of bp with a diffusion coefficient similar to that of other eukaryotic proteins known to encircle dsDNA ^31,33^. This diffusion contributed to targeting efficiently the fork position, which represents ∼0.16% of the total length of the scanned DNA. This ability would confer to the helicase an alternative mechanism for fork recognition within the ∼16 kbp long mtDNA. Twinkle remained stably bound at the fork, likely due to its interactions with the two strands of fork structure, as our data shows that ssDNA does not stall the helicase diffusion. Whether fast diffusion on dsDNA facilitates ATP-independent Twinkle activities, such as annealing and strand exchange, remains to be elucidated.

In addition, we collected evidence arguing that functional loading (i.e. loading followed by DNA unwinding initiation) is regulated by the ZBD and C-terminal tail of the helicase, together with the mtSSB. On the one hand, in the absence of mtSSB, eliminating the amino terminal ZBD favored detecting ΔZBD variant’s activities at tension 3 pN lower than that required to detect WT activity. This result suggest that the propensity of the ZBD to interact with DNA would interfere with the functional loading of the WT helicase on an SSB-free DNA fork. On the other hand, the presence of the mtSSB favored detection the WT helicase’s activities at significant lower tensions. This finding suggest that mtSSB binding to the DNA would outcompete the restrictive ZBD-DNA interactions that interfere with the functional loading of the WT helicase on the fork. Interestingly, this mtSSB stimulation was not observed for the ΔC-tail Twinkle variant lacking the unstructured C-terminal tail. The C-terminal tail of the homologous T7 helicase mediates physical protein-protein interactions^51^. Therefore, our results suggest that physical interactions between C-terminal tail of the helicase and mtSSB could be also relevant to promote functional loading of the helicase at the DNA fork. In line with these observations, it has been recently speculated that mtSSB may affect the way Twinkle loads on DNA^26^.

Upon functional loading, Twinkle unwound DNA with average rates (0.7-4 bp/ s at 1-10 mM ATP, respectively, F= 5 pN) in line with that estimated previously for this helicase in air-AFM studies^26^ and similar to those of other AAA+ helicases^52,53^. The average DNA unwinding rate of Twinkle was dominated by the high probability of occupancy of transient state(s) that do not support DNA unwinding (considered as pauses in our work). Similarly, frequent pause states have been shown to modulate the average DNA unwinding rates of other prokaryotic and the eukaryotic replicative helicases^43,53^. However, the origin of pauses during DNA unwinding is poorly understood.

Our results showed that truncation of the ZBD or C-terminal tail from Twinkle decreased significantly the pause occupancy of these variants compared to the WT at the lowest ATP concentrations (<4 mM) and tensions (F<6pN) used in this study. These findings support the implication of these domains in regulating the DNA unwinding kinetics of the helicase. On the one hand, the differences in response to mechanical tension between the real-time kinetics of WT and ΔZBD helicases suggest that interactions between ZBD and ssDNA favor pause occupancy at low ATP concentrations and mechanical tensions. One possible explanation is that the flexible and dynamic nature of the N-terminal domain of Twinkle^13,27^, where the ZBD is located, may facilitate regulatory interactions with the two single-stranded DNA strands at the replication fork. These interactions would be diminished by application of mechanical tension explaining the decrease of pause occupancy of WT with increasing tension.

On the other hand, the pause occupancy of WT and ΔZBD depend strongly on ATP concentration, whereas the ΔC-tail variant with enhanced ATPase activity shows no such dependency. These findings, together with the relatively high Km value of Twinkle for ATP (∼1.4 mM)^4,8^, point to ATP binding or exchange as a limiting factor and the C-terminal tail of the helicase as a relevant regulatory region that modulates ATP turnover and influences pause occupancy and DNA unwinding kinetics. Notably, the alphafold3 prediction of human Twinkle in the presence of ATP, which matches the structure of LcTwinkle bound to DNA and ATP^27^, shows the C-tail of Twinkle covering the ATP binding pocket and stabilizing the nucleotide (Figure S10). The basic C-tail may function as an auto-inhibitory lid, modulating ATP binding and exchange. This in turn, would hinder the proper coordination between ATP turnover and DNA binding by ZBD needed to couple translocation with DNA unwinding at near physiological ATP concentrations. Similarly, auto inhibitory basic C-terminal domains have been reported to hamper the activity of repair helicases^54^.

Autoregulation of the DNA unwinding rate would prevent the helicase from surpassing the DNA polymerase during occasional decoupling of the two enzymes at the replication fork. However, this auto-regulation should be released by the other replisome partners during active DNA replication. Our results showed that mtSSB stimulates DNA unwinding rate up to ∼3-4-fold by decreasing the pause occupancy of the helicase. The stimulation was specific to mtSSB (equal concentrations of EcoSSB did not stimulate, in agreement with previous bulk studies^24^), and decreased rapidly with increasing mechanical tension. These results indicated that stimulation of the DNA unwinding rate: 1) Is not due to facilitated fraying of the fork and ‘passive’ coverage of the newly generated ssDNA by the SSBs and, 2) depends on the tension-sensitive mtSSB binding mode^48^. We note that the real-time kinetics of the WT helicase in the presence of mtSSB was identical to that of ΔZBD variant in the absence of mtSSB at the lowest tension (2 pN). This observation suggests that mtSSB may outcompete specifically ZBD for DNA binding, as expected during functional loading of the helicase at the fork, releasing the auto-regulatory effect of this domain on the DNA unwinding kinetics.

The activity of WT Twinkle stalled after unwinding ∼40 bp of the fork. Remarkably, truncation of the ZBD stimulated the average number of unwound nucleotides up to ∼3 times. These results indicate that ZBD-DNA interactions play a role in controlling helicase processivity. Together with previous studies suggesting that Twinkle establishes multiple contacts with the displaced strand on its surface^3,5,6,19^, our findings support a scenario where increasing interactions of the ZBD and helicase surface with the two strands of the fork could build up torsional stress as DNA unwinding proceeds. This stress would eventually decouple ATPase activity and/or DNA binding from motor function, thus limiting processivity. The fact that the unwinding processivity decreased with increasing tension suggests that the experimental pulling geometry would contribute to stress generation at the fork. Remarkably, binding of mtSSB to the DNA stimulated up to ∼3 times the average processivity of the WT, but not that of the ΔZBD variant. These results strongly suggest that mtSSB would outcompete ZBD for DNA interactions, releasing the restrictive effect of this domain and enhancing the processivity of the helicase at the lowest tension (2 pN).

Upon stalling, Twinkle remained stably bound at the fork and prevented fast spontaneous closure of the hairpin. Instead, partial rewinding events were observed upon unwinding at <10 mM ATP. Under these conditions, the average number of rewound base pairs by the native helicase was ∼20, which contrasts with the average unwinding processivity of ∼40 bp. This difference implies that rewinding stops 20 nucleotides before the primer-template junction. This length coincides with the expected Twinkle DNA-binding size^4^ suggesting that during rewinding, the helicase could track the displaced strand until encountering the dsDNA primer-template junction. The different effects of tension, ATP and mtSSB on the rewinding probabilities and kinetics of the WT and helicase variants supports the following rewinding mechanism. Upon stalling, helicase interactions with the two strands of the fork and ATP turnover would favor a helicase-DNA conformation prone to move back gradually towards the primer-template junction pushed by the regression pressure of the fork. The ZBD and C-tail regions of Twinkle would have contrasting and likely complementary, roles in regulating its rewinding activity. The ZBD-DNA interactions would promote rewinding, while autoregulation of the ATPase activity by the C-tail would hinder this reaction. The presence of mtSSB would compete with the helicase for DNA interactions inhibiting rewinding at near physiological ATP concentrations. The requirements for rewinding resemble those described for the ‘intermolecular’ DNA annealing activity of Twinkle^4^. However, these two processes may be mechanistically different.

In summary, our findings indicate that Twinkle activity is strongly auto-regulated by its amino and carboxyl terminal domains. ZBD-DNA interactions and the control of ATP hydrolysis by the C-terminal tail serve as crucial regulatory checkpoints that modulate functional loading, DNA unwinding kinetics, and rewinding activities of the helicase at the fork. This auto-regulation is only partially alleviated by the mtSSB. Our work paves the way for future *in singulo* studies to explore the real-time kinetics of Twinkle, and its regulation, within the minimal human mitochondrial replisome during mtDNA replication.

### Methods Recombinant proteins

Recombinant wild-type (WT, i.e. 43-684) and the N- (ΔZBD, i.e. Δ145) and C- (ΔC-tail, i.e. 43-633) truncation variants of the human mitochondrial DNA helicase Twinkle were prepared from S*f9* cells as described previously^12^.

Cloning and purification of eGFP-Twinkle. The Twinkle DNA sequence was synthesized (gBlockTM, IDT) and cloned into a pRK5 vector for mammalian expression with a C-terminal eGFP-3xFLAG-6xHIS tag and a 3C-Prescission protease site using IVA cloning ^55^. Expi293FTM (ThermoFisher Scientific) cultured in Expi293FTM Expression Medium (ThermoFisher Scientific) at 3×106 cells/mL density were transfected with 1ng/ml of plasmid and 6ng/ml PeiMax® (Linear Polyethylenimine Hydrochloride: Polysciences) as a transfection reagent after 30 minutes of incubation at RT. Cells were harvested 48h after the transfection and frozen in liquid N_2_ for preservation and lysis. Frozen cells were resuspended in a ratio 1:3 (mass:volume) in Lysis buffer (25mM Hepes pH7.5, 500mM KCl, 10mM MgCl2, 5% glycerol, 2ug/ml Aprotinine, 5ug/ml Leupeptine, 1mM Benzamidine and 0.1 mg/ml AEBSF) and fully lysed by sonication in a Vibra-Cell 75042 sonicator (BioBlock Scientific) for 5 minutes, with pulses ON/OFF of 3 seconds ON-2 second OFF at 37% amplitude in an ice bath. Lysate was then cleared by centrifugation at 50.000G for 1h and filtered using a 0.22um syringe filter. ANTI-FLAG M2 Affinity beads (Sigma-Aldrich) were added to the cleared lysate and incubated for at least 3h at 4º in rotation. After the binding step, the beads were washed three times using the lysis buffer and transferred to a SigmaPrepTM spin column (Sigma-Aldrich in which we incubated the beads with the washing buffer with 300ng/uL of 3xFlag peptide (ChinaPeptide). After 30 minutes, the elution were recovered by centrifugation and frozen in liquid N_2_.

The eGFP-Polγ holoenzyme was prepared as described previously^56^ (see Figure S3 for more details).

Recombinant mtSSB was prepared from E. coli as described previously^57^. Recombinant EcoSSB was purchased from Thermofisher.

### Bulk biochemical assays

Biochemical assays are described in detail in the Supplementary information section (Figure S6).

### DNA constructs for single molecule experiments

Three DNA constructs were synthetized to quantify the diffusion on dsDNA and fork recognition by Twinkle-GFP. 1) Biotinylated dsDNA was generated from plasmid 89DIR.Poly-BbvCI (18709 bp)^58^. This plasmid was linearized with SalI-HF (NEB) and the resulting ends ligated to the complementary ends of biotinylated DNA handles amplified by PCR^58^. 2) dsDNA containing a single forked-like structure was generated upon linearization of vector 89DIR.Poly-BbvCI with SalI (NEB). Then specific locations between positions 6389 and 6452 bp of one strand of the plasmid were nicked with endonuclease Nt.BbvCI (NEB). The small DNA fragments between the nicked positions were thermally denatured by raising temperature to 80 °C rendering a 63 nt ssDNA gap. Then the two protruding regions ((dN)_30_) of a 446 bp long stem loop were hybridized and ligated overnight to the complementary ssDNA gap. Simultaneously, the plasmid ends were ligated to the complementary ends of biotinylated DNA fragments digested with XhoI-HF. Upon ligase denaturation, the DNA strand complementary to the section in which the hairpin was included (6389 and 6449 bp) was nicked at several positions with Nb.BbvCI (NEB). In the optical tweezers, individual DNA constructs tethered between streptavidin beads (see below) to induce the mechanical denaturation of the nicked region. This procedure generates a forked-liked structure with two ssDNA regions flanking a DNA hairpin. 3) Single-stranded DNA molecules (17303 nt) were obtained in the optical tweezers upon mechanical denaturation of a linear dsDNA, Figure S4. All protocols avoid DNA purification steps and enzymes were thermally denatured.

The circular dsDNA substrate with poly(dT)40 5’-tail for the HS-AFM experiments was constructed using pRC1765 mini plasmid (1765 bp,; Addgene plasmid # 141346). The mini plasmid was nicked at its two Nt.BbvCl sites and the resulting 33 nt stretch was replaced by an identical oligomer with poly(dT)_40_ extension at its 5’-end. The nick remaining at the 3’-end was ligated by the T4 DNA ligase.

The fork-like DNA construct for measuring the force and ATP-dependent unwinding kinetics of Twinkle was synthetize as described previously^48,59,60^. Briefly, the 3’- and 5’-ends of a 559 bp stem loop were ligated to a 2686 bp DNA handle labeled with digoxigenin and to a poly(dT)_35_ oligonucleotide functionalized with biotin, respectively. The poly(dT)_35_ tail facilitates loading of the helicase at the fork.

### Buffers

Single molecule studies were carried out in the reaction buffer containing 50 mM Tris pH 8.5, 30 mM KCl, 10 mM DTT, 4 or 10 mM MgCl_2_, 0.2 mg/ml BSA and ATP concentrations ranging from 1 mM to 10 mM.

### Single-molecule fluorescence imaging and data analysis

Single-molecule imaging experiments of Twinkle-GFP were performed on an instrument that combines three color confocal fluorescence microscopy (488, 532, 635 nm), with optical tweezers and microfluidics (C-trap® LUMICKS). Trap stiffness was adjusted to 0.32 pN/nm in both traps. DNA-molecules were trapped between two streptavidin-coated polystyrene beads (4.38 *µ*m diameter). The tethering of single DNA molecules was confirmed by analyzing the force-extension curve. The DNA was then transferred to another channel (channel 4) containing a 20 bp DNA oligo (50 nM) complementary to the 3’-tail of the forked structure and labelled with Atto 647N. Next, the DNA was transfer to channel 5 (previously passivated with BSA (2mg/ ml)), the distance between both beads was fixed to achieve a tension of 3 or 10 pN, and then Twinkle-eGFP (1 nM as hexamer) was flown into the channel. We used the 488 nm excitation laser for visualization of Twinkle-eGFP (emission filter of 500-525 nm) and the 635 nm excitation laser for visualization of the fork position labeled with Atto 647N (emission filter 650-750 nm). For confocal images (scans) the confocal pixel size was set to 100 nm and a scan velocity of 1 *µ*m/s^-1^. Kymographs were generated by single line scans covering the entire DNA stretched between the two beads and the edge of both beads. Pixel size was set to 100 nm and the illumination time per pixel was 0.1 ms, resulting in a typical time per line of 28.1 ms.

#### Software and code

We used Python 3.8 with several libraries for image processing: numpy==1.26.2, lumicks.pylake==1.0.0, matplotilb==3.7.1, scipy==1.10.1, pandas==2.0.1.

#### Trace tracking

We used the greedy algorithm^61–63^ implemented in Pylake Python library (https://github.com/lumicks/pylake//DOI10.5281/zenodo.4280788) to track the position (µm) of individual fluorescent spots with time (seconds). Kymograms were aligned using the position of the beads as fiducial points. The distance in µm was converted to base pairs (bp) using the theoretical average extension per bp according to the Worm Like Chain model. Similarly, we used the Freely Jointed Chain model of polymer elasticity to convert the distance in µm to single-stranded nucleotides. We used the *interp1* MATLAB function to interpolate data in time and in position.

#### Determination of number of fluorophores per diffraction-limited spot

The fluorescence background was quantified and subtracted from all scan images and kymograms. We used a eGFP-labelled version of the human mitochondrial polymerase holoenzyme (eGFP-Polγ) as a control to estimate the average number of eGFP-molecules per Twinkle diffraction–limited spot. Polγ holoenzyme is a heterotrimeric complex, consisting of the catalytic PolγA subunit and two PolγB accessory processivity factors. We labelled the PolγB subunit with eGFP. The reconstituted eGFP-Polγ holoenzyme, which binds stably to the primer-template position of the fork, provides an internal control to determine the intensity of two eGFP molecules. This intensity value was used to estimate the average number of eGFP-subunits per Twinkle diffraction–limited spots assuming a linear increase of the intensity with the number of eGFP molecules. First, we fit the first 50 maximum intensity data points of eGFP-Polγ diffraction–limited spots to a normal distribution to calculate the average intensity (*I*_*Pol*γ_) and standard deviation (σ_Polγ_) of two eGFPs 3.692 ± 0.788 intensity units (N=19 independent molecules). Then, an identical procedure was used to determine the average intensity of eGFP-Twinkle (*I*_*Tw*_) diffraction limited spots and its associated standard deviation (σ_Tw_). As the fluorophore intensity increases linearly with the number of eGPF per spot, upon determining the intensity of two eGFPs, we calculated the number of eGFP-labeled per Twinkle spot. The associated measurement error has the form:

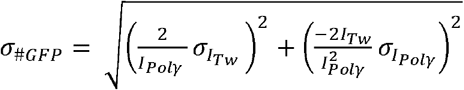

#### Diffusion analysis

We used the *DSigma2_Est* MATLAB function (https://www.nist.gov/programs-projects/fluctuations-and-nanoscale-control-software-archived) to estimate the Mean Squared Displacement (MSD) *vs* time lag (τ) for each trace as: 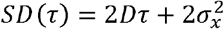, where *D* is the diffusion coefficient and *σ*_*x*_ is the location measurement error. The appropriate range of delay times were calculated as previously published^62,64^.

### HS-AFM imaging and data analysis

Circular DNA (1 nM) diluted in a buffer containing 50 mM Tris pH 8.5, 10 mM MgCl_2_ and 40 mM KCl was deposited on freshly cleaved bare mica. Upon excess DNA removal, the DNA was incubated for 5 minutes with Twinkle diluted to 1 nM in 40 *μ*l of the reaction buffer supplemented with 4 mM ATP. Then, we used the tapping mode of a custom-built HS-AFM apparatus ^65,66^ to scan an area of 300 × 300 nm^2^ (100×100 pixels) at 1-2 frames per second at room temperature (22 °C). The free-oscillation amplitude of the cantilever (A0) was set to ∼2 nm, and the set point of the feedback amplitude was set to ∼0.9 × Å. Small cantilevers (BL-AC10DS-A2Olympus) with a spring constant of ∼0.15 N/m, resonance frequency of ∼500 kHz, and quality factor of ∼1.5 in water were used. An EBD tip was grown for 1.5 min on the original tip of cantilever. Video frames were analyzed using custom-made algorithms implemented in IGOR Pro (WaveMetrics, USA).

### Single-molecule force spectroscopy experiments and data analysis

A counter propagating dual-beam optical tweezers instrument^34^ was used to manipulate individual DNA constructs functionalized with biotin or digoxigenin at each end. The DNA was tethered between a streptavidin- and anti-digoxigenin-coated polystyrene beads (3 *μ*m) held on top of a micropipette and in the optical trap, respectively. Twinkle (WT and variants) and SSB proteins were diluted to 5 nM in the reaction buffer and flown inside the flow cell. We note that mtSSB concentrations higher than 5 nM prevented detection of single-molecule DNA unwinding activities. This finding is in line with the deleterious effect of high mtSSB concentrations on the *in vitro* activity of Twinkle^18,24^. Data was monitored at 5100 Hz at 22±1ºC using a feedback loop to maintain a constant mechanical tension on the DNA construct below 13 picoNewtons (pN). The trap stiffness was *k*=0.135±0.0043 pN nm^-1^ (3.0 *μ*m beads).

The number of unwound base pairs (processivity) was obtained by dividing the increase of the tether extension (Δx) by the change in extension at a given tension accompanying during each catalytic step the generation of two new ssDNA nucleotides (or SSB-bound nucleotides). We used the average extensions per nucleotide of free- and SSB-bound ssDNA as a function of tension reported previously^48^.

The average DNA unwinding rate was determined by a line fit to the traces showing the number of replicated nucleotides versus time. The final average rate at each mechanical tension was obtained by averaging over all of the traces taken at that tension. The average replication rate without pauses or pause free velocity of each trace was determined as described previously^40,67^. Briefly, individual trajectories were analyzed with an algorithm that computes the instantaneous velocities, averaging the position over sliding time windows.

The pause occupancy represents the probability of finding the helicase in a transient, not active or pause state. The pause occupancy at each tension (F) was calculated as *Pocc*(*F*) = (1 – *Vmean*(*F*)/*Vmax*(*F*)), where *Vmean*(*F*) and *Vmax*(*F*) represent the average and pause-free replication rates at each tension, respectively.

## Acknowledgments

We are grateful to Prof. Laurie S. Kaguni for generous gift of expression constructs for Twinkle WT, and ΔZBD and ΔC-tail variants. We thank Cameron Young for helping with preparation of the figures. G.L.C. We thank the members of our labs for useful discussions.

## References

1. Spelbrink, J. N. et al. Human mitochondrial DNA deletions associated with mutations in the gene encoding Twinkle, a phage T7 gene 4-like protein localized in mitochondria. Nat. Genet. 28, 223–231 (2001).

2. Peter, B. & Falkenberg, M. TWINKLE and Other Human Mitochondrial DNA Helicases: Structure, Function and Disease. Genes 11, 408 (2020).

3. Sen, D., Patel, G. & Patel, S. S. Homologous DNA strand exchange activity of the human mitochondrial DNA helicase TWINKLE. Nucleic Acids Res. 44, 4200–4210 (2016).

4. Sen, D., Nandakumar, D.Tang, G.-Q. & Patel, S. S. Human Mitochondrial DNA Helicase TWINKLE Is Both an Unwinding and Annealing Helicase*. J. Biol. Chem. 287, 14545–14556 (2012).

5. Khan, I. et al. Biochemical Characterization of the Human Mitochondrial Replicative Twinkle Helicase: SUBSTRATE SPECIFICITY, DNA BRANCH MIGRATION, AND ABILITY TO OVERCOME BLOCKADES TO DNA UNWINDING *. J. Biol. Chem. 291, 14324–14339 (2016).

6. Singh, A., Patel, G. & Patel, S. S. Twinkle-Catalyzed Toehold-Mediated DNA Strand Displacement Reaction. J. Am. Chem. Soc. 145, 24522–24534 (2023).

7. Hensen, F. et al. Mitochondrial RNA granules are critically dependent on mtDNA replication factors Twinkle and mtSSB. Nucleic Acids Res. 47, 3680–3698 (2019).

8. Longley, M. J., Humble, M. M., Sharief, F. S. & Copeland, W. C. Disease variants of the human mitochondrial DNA helicase encoded by C10orf2 differentially alter protein stability, nucleotide hydrolysis, and helicase activity. J. Biol. Chem. 285, 29690–29702 (2010).

9. Matsushima, Y. & Kaguni, L. S. Differential Phenotypes of Active Site and Human Autosomal Dominant Progressive External Ophthalmoplegia Mutations in Drosophila Mitochondrial DNA Helicase Expressed in Schneider Cells*. J. Biol. Chem. 282, 9436–9444 (2007).

10. Tyynismaa, H. et al. Mutant mitochondrial helicase Twinkle causes multiple mtDNA deletions and a late-onset mitochondrial disease in mice. Proc. Natl. Acad. Sci. 102, 17687– 17692 (2005).

11. Tyynismaa, H. & Suomalainen, A. Mouse models of mitochondrial DNA defects and their relevance for human disease. EMBO Rep. 10, 137 (2009).

12. Ziebarth, T. D., Farr, C. L. & Kaguni, L. S. Modular Architecture of the Hexameric Human Mitochondrial DNA Helicase. J. Mol. Biol. 367, 1382–1391 (2007).

13. Fernández-Millán, P. et al. The hexameric structure of the human mitochondrial replicative helicase Twinkle. Nucleic Acids Res. 43, 4284–4295 (2015).

14. Riccio, A. A. et al. Structural insight and characterization of human Twinkle helicase in mitochondrial disease. Proc. Natl. Acad. Sci. 119, e2207459119 (2022).

15. Shutt, T. E. & Gray, M. W. Twinkle, the Mitochondrial Replicative DNA Helicase, Is Widespread in the Eukaryotic Radiation and May Also Be the Mitochondrial DNA Primase in Most Eukaryotes. J. Mol. Evol. 62, 588–599 (2006).

16. Singleton, M. R., Dillingham, M. S. & Wigley, D. B. Structure and Mechanism of Helicases and Nucleic Acid Translocases. Annu. Rev. Biochem. 76, 23–50 (2007).

17. Matsushima, Y., Farr, C. L., Fan, L. & Kaguni, L. S. Physiological and Biochemical Defects in Carboxyl-terminal Mutants of Mitochondrial DNA Helicase. J. Biol. Chem. 283, 23964–23971 (2008).

18. Rodrigues, A. P. C. & Oliveira, M. T. Stimulation of Variant Forms of the Mitochondrial DNA HelicaseDNA helicases Twinkle by the Mitochondrial Single-Stranded DNA-Binding ProteinMitochondrial single-stranded DNA-binding protein (mtSSB). in Single Stranded DNA Binding Proteins (ed. Oliveira, M. T.) 313–322 (Springer US, New York, NY, 2021). doi:10.1007/978-1-0716-1290-3_20.

19. Johnson, L. C., Singh, A. & Patel, S. S. The N-terminal domain of human mitochondrial helicase Twinkle has DNA-binding activity crucial for supporting processive DNA synthesis by polymerase γ. J. Biol. Chem. 299, 102797 (2023).

20. Farge, G. et al. The N-terminal domain of TWINKLE contributes to single-stranded DNA binding and DNA helicase activities. Nucleic Acids Res. 36, 393–403 (2008).

21. Lee, S.-J., Zhu, B., Akabayov, B. & Richardson, C. C. Zinc-binding Domain of the Bacteriophage T7 DNA Primase Modulates Binding to the DNA Template*. J. Biol. Chem. 287, 39030–39040 (2012).

22. Kaguni, L. S. & Oliveira, M. T. Structure, function and evolution of the animal mitochondrial replicative DNA helicase. Crit. Rev. Biochem. Mol. Biol. 51, 53–64 (2016).

23. Ziebarth, T. D. et al. Dynamic effects of cofactors and DNA on the oligomeric state of human mitochondrial DNA helicase. J. Biol. Chem. 285, 14639–14647 (2010).

24. Korhonen, J. A., Gaspari, M. & Falkenberg, M. TWINKLE Has 51⍰ → 31⍰ DNA Helicase Activity and Is Specifically Stimulated by Mitochondrial Single-stranded DNA-binding Protein*. J. Biol. Chem. 278, 48627–48632 (2003).

25. Jemt, E. et al. The mitochondrial DNA helicase TWINKLE can assemble on a closed circular template and support initiation of DNA synthesis. Nucleic Acids Res. 39, 9238–9249 (2011).

26. Kaur, P. et al. Single-molecule level structural dynamics of DNA unwinding by human mitochondrial Twinkle helicase. J. Biol. Chem. 295, 5564–5576 (2020).

27. Li, Z. et al. Structural and dynamic basis of DNA capture and translocation by mitochondrial Twinkle helicase. Nucleic Acids Res. 50, 11965–11978 (2022).

28. Korhonen, J. A., Pham, X. H., Pellegrini, M. & Falkenberg, M. Reconstitution of a minimal mtDNA replisome in vitro. EMBO J. 23, 2423–2429 (2004).

29. Carney, S. M. & Trakselis, M. A. The excluded DNA strand is SEW important for hexameric helicase unwinding. Methods 108, 79–91 (2016).

30. Candelli, A., Wuite, G. J. L. & Peterman, E. J. G. Combining optical trapping, fluorescence microscopy and micro-fluidics for single molecule studies of DNA–protein interactions. Phys. Chem. Chem. Phys. 13, 7263–7272 (2011).

31. Alcón, P. et al. FANCD2–FANCI surveys DNA and recognizes double-to single-stranded junctions. Nature 632, 1165–1173 (2024).

32. Sánchez, H. et al. DNA replication origins retain mobile licensing proteins. Nat. Commun. 12, 1908 (2021).

33. Wasserman, M. R., Schauer, G. D., O’Donnell, M. E. & Liu, S. Replication Fork Activation Is Enabled by a Single-Stranded DNA Gate in CMG Helicase. Cell 178, 600-611.e16 (2019).

34. Smith, S. B., Cui, Y. & Bustamante, C. [7] Optical-trap force transducer that operates by direct measurement of light momentum. in Methods in Enzymology vol. 361 134–162 (Academic Press, 2003).

35. Sun, B. et al. ATP-induced helicase slippage reveals highly coordinated subunits. Nature 478, 132–135 (2011).

36. Yuan, Z. et al. DNA unwinding mechanism of a eukaryotic replicative CMG helicase. Nat. Commun. 11, 688 (2020).

37. Spinks, R. R., Spenkelink, L. M., Dixon, N. E. & van Oijen, A. M. Single-Molecule Insights Into the Dynamics of Replicative Helicases. Front. Mol. Biosci. 8, (2021).

38. Bocanegra, R., Plaza, G. I., Pulido, C. R. & Ibarra, B. DNA replication machinery: Insights from in vitro single-molecule approaches. Comput. Struct. Biotechnol. J. 19, 2057 (2021).

39. Schlierf, M., Wang, G., Chen, X. S. & Ha, T. Hexameric helicase G40P unwinds DNA in single base pair steps. eLife 8, e42001 (2019).

40. Cerrón, F. et al. Replicative DNA polymerases promote active displacement of SSB proteins during lagging strand synthesis. Nucleic Acids Res. 47, 5723–5734 (2019).

41. Kennedy, H. J. et al. Glucose Generates Sub-plasma Membrane ATP Microdomains in Single Islet β-Cells: POTENTIAL ROLE FOR STRATEGICALLY LOCATED MITOCHONDRIA *. J. Biol. Chem. 274, 13281–13291 (1999).

42. Gajewski, C. D., Yang, L., Schon, E. A. & Manfredi, G. New Insights into the Bioenergetics of Mitochondrial Disorders Using Intracellular ATP Reporters. Mol. Biol. Cell 14, 3628–3635 (2003).

43. Ribeck, N., Kaplan, D. L., Bruck, I. & Saleh, O. A. DnaB Helicase Activity Is Modulated by DNA Geometry and Force. Biophys. J. 99, 2170–2179 (2010).

44. Ribeck, N. & Saleh, O. A. DNA Unwinding by Ring-Shaped T4 Helicase gp41 Is Hindered by Tension on the Occluded Strand. PLOS ONE 8, e79237 (2013).

45. Manosas, M., Spiering, M. M., Ding, F., Croquette, V. & Benkovic, S. J. Collaborative coupling between polymerase and helicase for leading-strand synthesis. Nucleic Acids Res. 40, 6187–6198 (2012).

46. Johnson, D. S., Bai, L., Smith, B. Y., Patel, S. S. & Wang, M. D. Single-Molecule Studies Reveal Dynamics of DNA Unwinding by the Ring-Shaped T7 Helicase. Cell 129, 1299–1309 (2007).

47. Suksombat, S., Khafizov, R., Kozlov, A. G., Lohman, T. M. & Chemla, Y. R. Structural dynamics of E. coli single-stranded DNA binding protein reveal DNA wrapping and unwrapping pathways. eLife 4, e08193 (2015).

48. Morin, J. A. et al. DNA synthesis determines the binding mode of the human mitochondrial single-stranded DNA-binding protein. Nucleic Acids Res. 45, 7237–7248 (2017).

49. O’Shea, V. L. & Berger, J. M. Loading strategies of ring-shaped nucleic acid translocases and helicases. Curr. Opin. Struct. Biol. 25, 16–24 (2014).

50. Bleichert, F., Botchan, M. R. & Berger, J. M. Mechanisms for initiating cellular DNA replication. Science 355, eaah6317 (2017).

51. Lee, S.-J. & Richardson, C. C. Choreography of bacteriophage T7 DNA replication. Curr. Opin. Chem. Biol. 15, 580–586 (2011).

52. Joo, S., Chung, B. H., Lee, M. & Ha, T. H. Ring-shaped replicative helicase encircles double-stranded DNA during unwinding. Nucleic Acids Res. 47, 11344–11354 (2019).

53. Burnham, D. R., Kose, H. B., Hoyle, R. B. & Yardimci, H. The mechanism of DNA unwinding by the eukaryotic replicative helicase. Nat. Commun. 10, 2159 (2019).

54. Richards, J. et al. STRUCTURE OF THE DNA REPAIR HELICASE HEL308 REVEALS DNA BINDING AND AUTOINHIBITORY DOMAINS. J. Biol. Chem. 283, 5118–5126 (2008).

55. García-Nafría, J., Watson, J. F. & Greger, I. H. IVA cloning: A single-tube universal cloning system exploiting bacterial In Vivo Assembly. Sci. Rep. 6, 27459 (2016).

56. Fernández-Moreno, M. A., Farr, C. L., Kaguni, L. S. & Garesse, R. Drosophila melanogaster as a Model System to Study Mitochondrial Biology. in Mitochondria: Practical Protocols (eds. Leister, D. & Herrmann, J.M.) 33–49 (Humana Press, Totowa, NJ, 2007). doi:10.1007/978-1-59745-365-3_3.

57. Ciesielski, G. L. et al. Mitochondrial Single-stranded DNA-binding Proteins Stimulate the Activity of DNA Polymerase γ by Organization of the Template DNA. J. Biol. Chem. 290, 28697–28707 (2015).

58. Aicart-Ramos, C., Hormeno, S., Wilkinson, O. J., Dillingham, M. S. & Moreno-Herrero, F. Chapter Twelve - Long DNA constructs to study helicases and nucleic acid translocases using optical tweezers. in Methods in Enzymology (ed. Trakselis, M. A.) vol. 673 311–358 (Academic Press, 2022).

59. Morin, J. A. et al. Active DNA unwinding dynamics during processive DNA replication. Proc. Natl. Acad. Sci. 109, 8115–8120 (2012).

60. Morin, J. A. et al. Manipulation of single polymerase-DNA complexes: a mechanical view of DNA unwinding during replication. Cell Cycle Georget. Tex 11, 2967–2968 (2012).

61. Berglund, A. J., McMahon, M. D., McClelland, J. J. & Liddle, J. A. Fast, bias-free algorithm for tracking single particles with variable size and shape. Opt. Express 16, 14064– 14075 (2008).

62. Berglund, A. J. Statistics of camera-based single-particle tracking. Phys. Rev. E 82, 011917 (2010).

63. Sbalzarini, I. F. & Koumoutsakos, P. Feature point tracking and trajectory analysis for video imaging in cell biology. J. Struct. Biol. 151, 182–195 (2005).

64. Michalet, X. & Berglund, A. J. Optimal diffusion coefficient estimation in single-particle tracking. Phys. Rev. E Stat. Nonlin. Soft Matter Phys. 85, 061916 (2012).

65. Fukuda, S. & Ando, T. Technical advances in high-speed atomic force microscopy. Biophys. Rev. 15, 2045–2058 (2023).

66. Ando, T., Uchihashi, T. & Fukuma, T. High-speed atomic force microscopy for nano-visualization of dynamic biomolecular processes. Prog. Surf. Sci. 83, 337–437 (2008).

67. Plaza-G A I., et al. Mechanism of strand displacement DNA synthesis by the coordinated activities of human mitochondrial DNA polymerase and SSB. Nucleic Acids Res. 51, 1750–1765 (2023).

68. Jumper, J. et al. Highly accurate protein structure prediction with AlphaFold. Nature 596, 583–589 (2021).

