## Supplemental Information for "Auto-regulation of the real-time kinetics of the human mitochondrial replicative helicase"

^1^Instituto Madrileño de Estudios Avanzados en Nanociencia, IMDEA Nanociencia, Faraday 9, 28049 Madrid, Spain, ^2^Department of Biology, University of North Florida, Jacksonville, FL, United States. ^3^Structural Biology Programme, Spanish National Cancer Research Centre (CNIO), Madrid, Spain. ^4^Centro Nacional de Biotecnología Campus de Cantoblanco, 28049, Madrid, Spain. ^5^Nanobiotecnología (IMDEA-Nanociencia), Unidad Asociada al Centro Nacional de Biotecnología (CSIC), 28049 Madrid, Spain

**^†^** Current address: Quantemol Ltd, London, England, UK.

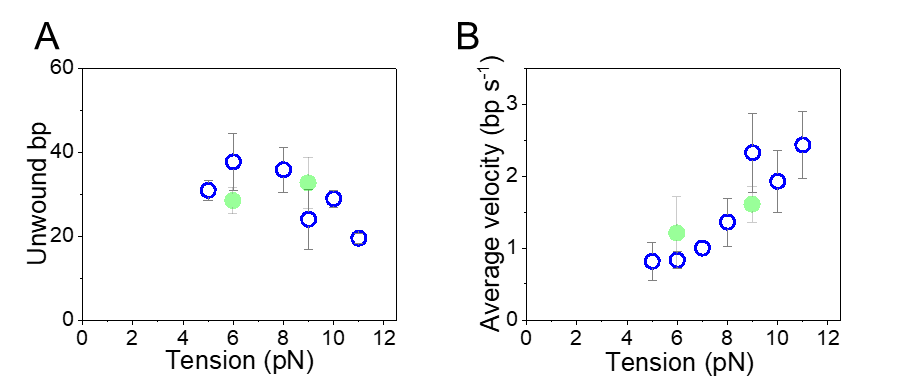

**Figure S1: eGFP-Twinkle presents identical DNA unwinding kinetics than WT Twinkle**. Tension dependencies of the average unwinding processivities (**A**) and average unwinding rate (**B**) of WT Twinkle (blue symbols) and eGFP-Twinkle (green symbols). For the two data plots data was taken at 4 mM ATP and error bars represent standard error of the mean (s.e.).

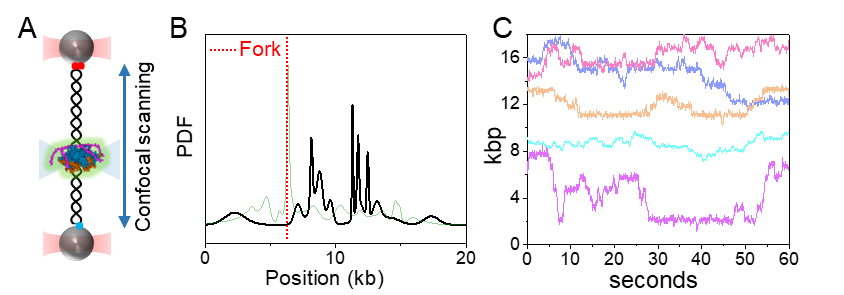

**Figure S2:** **Localization and** **diffusion of eGFP-Twinkle on dsDNA without fork. A)** Diagram of the optical tweezers-confocal assay to image eGFP-Twinkle on individual dsDNA molecules attached between optically trapped beads. **B)** Distribution of initial positions of eGFP-Twinkle diffraction-limited spots on dsDNA molecules lacking the DNA fork (black, N= 30). eGFP-Twinkle did not present preferential binding positions on dsDNA. For comparison, the initial distribution of eGFP-Twinkle on dsDNA containing a fork is shown in pale green (as in Figure 2C in the main text). Red dotted line indicates the position of the fork. **C)** Representative position vs. time plots of eGFP-Twinkle spots along dsDNA without a forked structure (4mM ATP). Diffusion of eGFP-Twinkle was not halted at any particular position on fork-free dsDNA, which contrast with results on dsDNA constructs bearing a DNA fork (Figure 2E). Traces were shifted along the Y-axis for clarity of display.

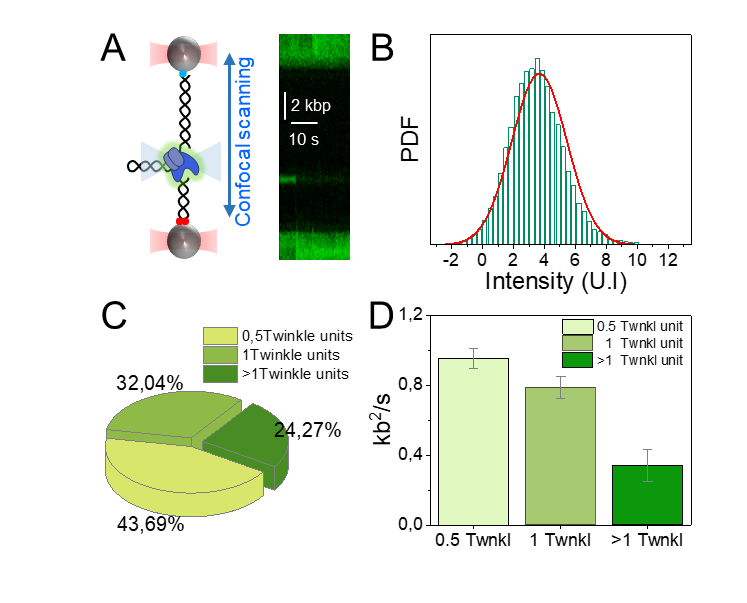

**Figure S3: Determination of number of fluorophores per diffraction-limited spot**. We used a human mitochondrial polymerase labelled with two eGFP (eGFP-Polγ) as a control to estimate the average number of eGFP-molecules per Twinkle diffraction–limited spot. eGFP-Polγ holoenzyme was prepared by combining PolγA (Ciesielski Lab collection) with eGFP-PolγB conjugate at 1:2 molar ratio (i.e., 1:1 PolγA : eGFP-PolγB dimer). The eGFP-PolγB coding sequence was synthesized and subcloned to pET21a(+) NdeI/HindIII site, by GenScript. The sequence includes N-terminal his-tag linked to eGFP sequence via DPPV linker, followed by SGLRSRA linker and PolγB Δ25 sequence, which is similar to previously reported PolγB-GFP fusions^1^. The fusion protein was purified as reported previously^2^. **A)** (Left) Schematic representation of the experimental setup showing the reconstituted eGFP-Polγ holoenzyme (blue-green) bound to the primer-template position of a DNA fork held between two optically trapped beads (as described in the main text). (Right) Representative kymogram showing stable binding of eGFP-Polγ (green) at the forked DNA position and subsequent photo bleaching. **B)** The distribution intensity of the initial 50 data points of eGFP-Polγ diffraction–limited spots (N=19) was fit to a normal distribution to calculate the average intensity and standard deviation of two eGFPs, 3.692 ± 0.788. An identical procedure was used to determine the average intensity of eGFP-Twinkle diffraction limited spots and its associated standard deviation. The number of eGFP-labeled per Twinkle spot was then estimated assuming a linear relationship between the fluorophore intensity with the number of eGPF per spot. **C)** Distribution of Twinkle oligomeric states along dsDNA. Twinkle binds dsDNA in different oligomeric states including broken rings (~44%), Twinkle units (~32%) and higher oligomers (~24%). The distribution of oligomeric states along the dsDNA was in line to that found by previous structural studies for human Twinkle W315L variant in solution^3^. Therefore, these results show that Twinkle has the ability to bind dsDNA in different oligomeric states. **D)** Oligomeric assemblies containing 6.0±2 eGFPs were considered as individual Twinkle units (1 Twnkl) and multiples of this value as either partial (0.5 Twnkl) or higher oligomeric forms (>1 Twnkl). The plot shows the diffusion coefficients on dsDNA of each oligomeric state. Error bars show s.e.

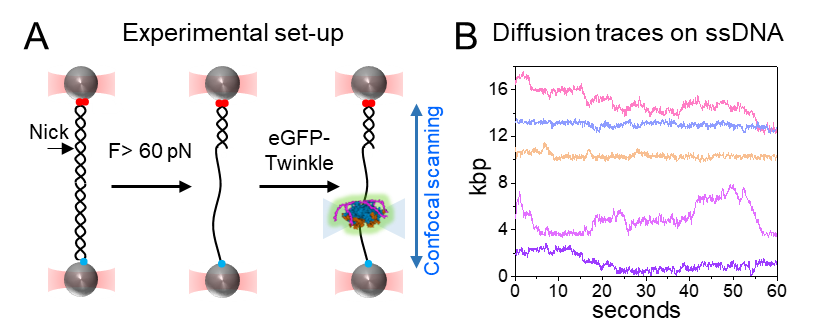

**Figure S4. Diffusion traces of eGFP-Twinkle on ssDNA. A)** Schematic of the experimental setup. ssDNA was generated by mechanical denaturation of dsDNA as described previously^4^. Briefly, a dsDNA molecule (18,709 bp) containing a single nick at one strand was stretched between two polystyrene beads held by two optical traps. Force (F) above 60 pN induces the mechanical denaturation of the molecule and the release of the nicked strand. This resulted in a DNA molecule containing a stretch of ssDNA of 17,303 nt. The DNA is then transfer to a flow channel containing eGFP-Twinkle diluted to 5 nM in the reaction buffer containing 4 mM ATP (Methods) and was held at constant tension of 10 pN. **B)** Representative diffusion traces of eGFP-Twinkle units on ssDNA. Traces were shifted along the Y-axis for clarity of display.

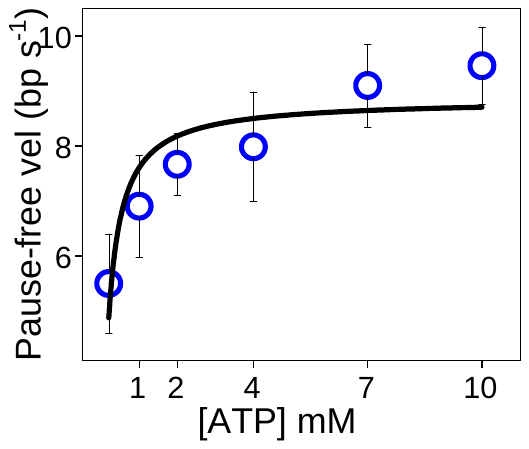

**Figure S5**: **The pause-free velocity of the WT Twinkle shows a sigmoidal dependency with ATP concentration**. The black line represents the fit to the data using the Michaelis-Menten expression. Error bars represent s.e.

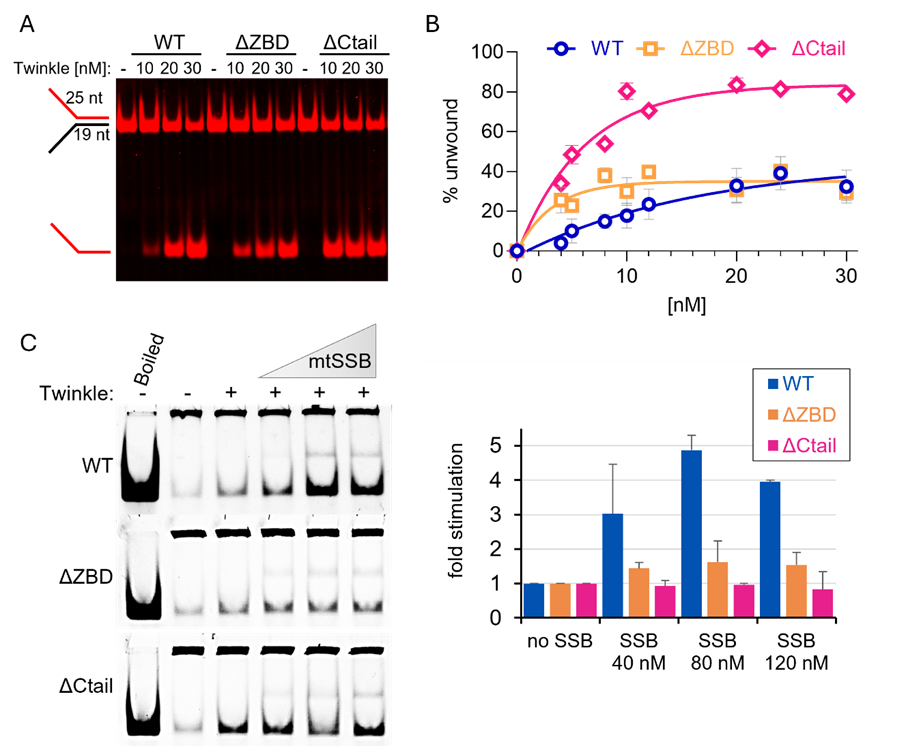

**Figure S6: Biochemical analysis of the unwinding activity of Twinkle WT, ΔZBD, and ΔC-tail variants**. **A)** Representative image of the helicase unwinding assay. Reaction was conducted as described below, using 2 nM fluorescently labeled DNA fork (depicted on the side) and indicated concentrations of Twinkle variants. **B)** Quantitative analysis of the DNA fork unwinding by the Twinkle variants. Reactions were conducted as described below, using 0.5 nM radioactively labeled DNA fork and indicated increments of Twinkle variants. The product abundance was normalized to total DNA substrate. The data represents an average of at least three independent experiments. Error bars show standard deviation. **C)** The mtSSB stimulation assay was conducted as described below. The panel on the left demonstrates representative gel images, and the graph on the right represents quantitation of at least three independent experiment repeats. The product abundance was normalized to total DNA substrate and the fold stimulation was calculated as fold increase over the product generated without mtSSB addition. Error bars show standard deviation.

**Helicase unwinding assay** was performed as described previously^5^, using fork DNA substrate 1 (25 nt 5’ and 3’ tails followed by 19 nt duplex) and Twinkle variants under study at indicated concentrations. Oligonucleotides used to assemble the DNA substrate were purchased from IDT. The 5’-tail was either labeled fluorescently (Cy5) or radioactively with ^32^P, as indicated. Reaction conditions were consistent with single molecule experiments, i.e., 50 mM Tris pH 8.5, 30 mM KCl, 10 mM DTT, 4 mM MgCl_2_, 0.1 mg/ml BSA, 10% glycerol and 5 mM ATP, for 30 min. at 37°C. The products were separated by electrophoresis through a 12% non-denaturing polyacrylamide gel. When fluorescently labeled DNA substrate was used, products were visualized directly on the gel. When radioactive label was used, gels were dried onto DE81 (Whatman) and auto-radiographed overnight at 80 °C with an intensifying screen. Intensities of the bands were quantified by densitometry using ImageJ (NIH). The obtained values were plotted and fitted to the single-phase association trendline in GraphPad Prism.

**mtSSB stimulation assay** was based on previously published approaches^6,7^. Briefly, reaction mixture contained 2 nM single-stranded circular M13mp18 DNA (New England Biolabs) primed with 5’Cy5-labeled oligonucleotide ACATGATAAGATACATGGATGAGTTTGGACAAACCACAACGTAAAACGACGGCCAGTGCC (IDT), yielding a 20 nt duplex segment with 40 nt 5’ tail^7^. The reaction buffer included 50 mM Tris pH 8.5, 30 mM KCl, 10 mM DTT, 4 mM MgCl_2_, 0.1 mg/ml BSA, 10% glycerol, 2nM Twinkle variant, and indicated concentrations of mtSSB. Unwinding reactions were initiated by the addition of 5 mM ATP, and carried out for 30 minutes at 37°C. Reaction was stopped with 10x stop solution: 100 mM EDTA, 6% SDS, 50% glycerol. We did not observe any significant levels of spontaneous reannealing of unwound DNA under the applied assay conditions. Products were next analyzed on native 10 % polyacrylamide (49.5:0.5) gel and scanned for Cy5 fluorescence. Lane labeled as ‘Boiled’ represents a DNA substrate sample incubated at 98°C for 5 minutes immediately before loading. The relative abundance of the unwinding product was quantified in ImageJ (NIH).

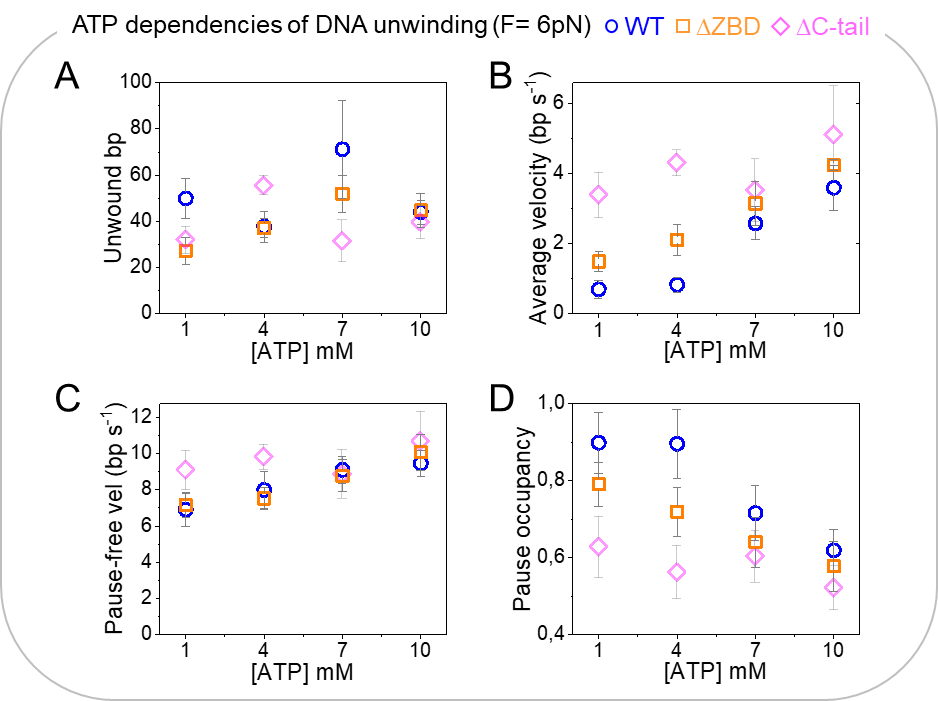

**Figure S7: The DNA unwinding kinetics of ΔZBD variant depend on ATP concentration in a manner similar to that of the WT helicase**. ATP dependencies of the average unwinding processivity (**A**), average unwinding rate (**B**), pause-free velocity (**C**) and pause occupancy (**E**) for the ΔZBD variant are indicated by orange symbols. For comparison, we included WT and ΔC-tail variant data in blue and magenta, respectively. Data was recorded at F=6 pN, which is the minimum tension that consistently allowed detection of activities at all ATP concentrations. Error bars in all data plots represent s.e.

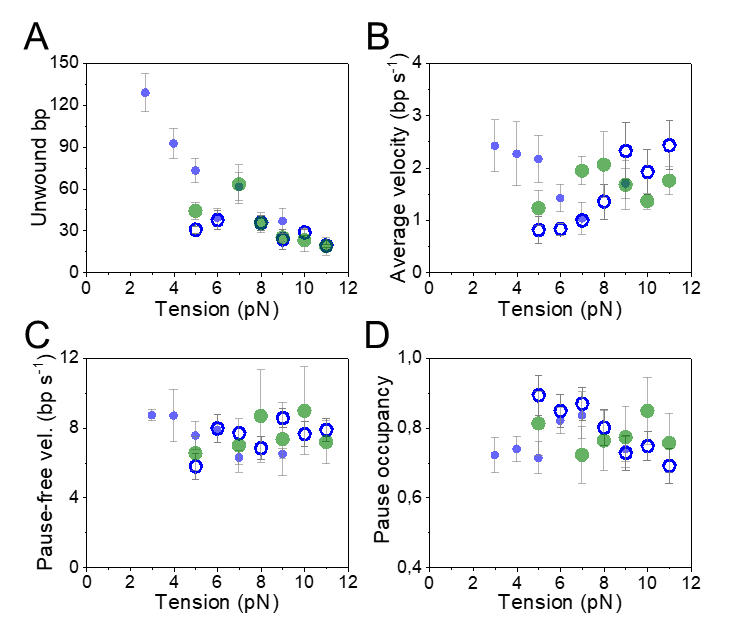

**Figure S8**: ***E. coli* SSB does not stimulate the real-time DNA unwinding kinetics of Twinkle.** Tension dependencies of the average unwinding processivity (**A**), average unwinding rate (**B**), pause-free velocity (**C**) and pause occupancy (**D**) for the WT helicase in the absence (blue empty symbols) and presence of 5 mM mtSSB (solid blue symbols) or 5 mM EcoSSB (green symbols). Data was collected at 4 mM ATP. For all plots error bars represent s.e.

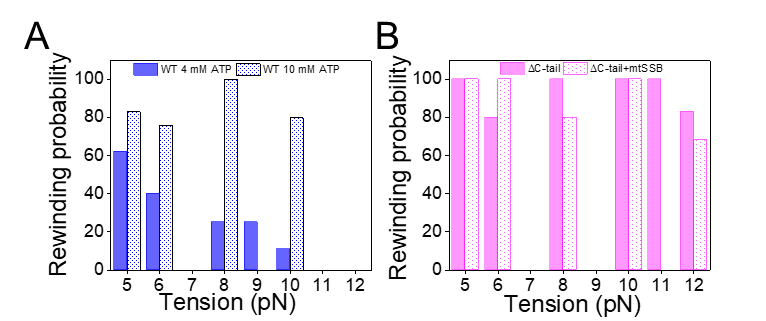

**Figure S9**: **Probabilities of occurrence of rewinding events upon DNA unwinding stalling as a function of tension**. **A)** At 4 mM ATP, the rewinding probability of WT Twinkle decreased with increasing tension (blue columns). In contrast, the rewinding probability was tension independent at 10 mM ATP (dotted columns). **B)** mtSSB (5 nM) did not inhibited the rewinding probability of the ΔC-tail variant at any tension. These results highlight that elevated ATP turnover would favor rewinding events even at high tension and/or in the presence of mtSBB bound to the unwound DNA strands.

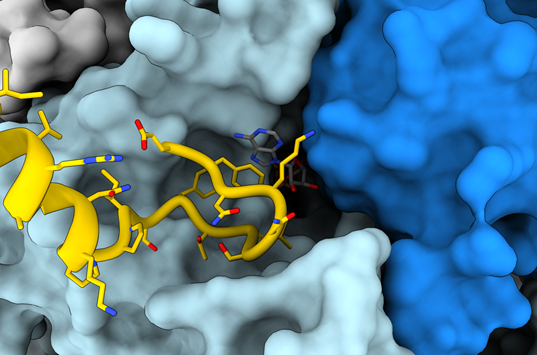

**Figure S10**: **Alphafold3 model of human Twinkle ATP binding pocket.** Surface representation of adjacent monomers (in light and dark blue) with ATP in the active site of the human Twinkle hexamer. Residues 620-684 of the light blue-monomer (in yellow) are displayed in cartoon and stick representation to highlight their interaction with the ATP (Phe621) and their potential role as an active site lid. The alphafold3^8^ model of the human Twinkle hexamer with ATP closely resembles the LcTwinkle structure with DNA and ATP^9^.

**Supplementary references:**

1.	Young, M. J., Humble, M. M., DeBalsi, K. L., Sun, K. Y. & Copeland, W. C. POLG2 disease variants: analyses reveal a dominant negative heterodimer, altered mitochondrial localization and impaired respiratory capacity. *Hum. Mol. Genet.* **24**, 5184 (2015).

2.	Oliveira, M. T. & Kaguni, L. S. Comparative purification strategies for Drosophila and human mitochondrial DNA replication proteins: DNA polymerase gamma and mitochondrial single-stranded DNA-binding protein. *Methods Mol. Biol. Clifton NJ* **554**, 37–58 (2009).

3.	Riccio, A. A. *et al.* Structural insight and characterization of human Twinkle helicase in mitochondrial disease. *Proc. Natl. Acad. Sci.* **119**, e2207459119 (2022).

4.	Aicart-Ramos, C., Hormeno, S., Wilkinson, O. J., Dillingham, M. S. & Moreno-Herrero, F. Chapter Twelve - Long DNA constructs to study helicases and nucleic acid translocases using optical tweezers. in *Methods in Enzymology* (ed. Trakselis, M. A.) vol. 673 311–358 (Academic Press, 2022).

5.	Khan, I. *et al.* Biochemical Characterization of the Human Mitochondrial Replicative Twinkle Helicase. *J. Biol. Chem.* **291**, 14324–14339 (2016).

6.	Rodrigues, A. P. C. & Oliveira, M. T. Stimulation of Variant Forms of the Mitochondrial DNA HelicaseDNA helicases Twinkle by the Mitochondrial Single-Stranded DNA-Binding ProteinMitochondrial single-stranded DNA-binding protein (mtSSB). in *Single Stranded DNA Binding Proteins* (ed. Oliveira, M. T.) 313–322 (Springer US, New York, NY, 2021). doi:10.1007/978-1-0716-1290-3_20.

7.	Korhonen, J. A., Gaspari, M. & Falkenberg, M. TWINKLE Has 5′ → 3′ DNA Helicase Activity and Is Specifically Stimulated by Mitochondrial Single-stranded DNA-binding Protein*. *J. Biol. Chem.* **278**, 48627–48632 (2003).

8.	Jumper, J. *et al.* Highly accurate protein structure prediction with AlphaFold. *Nature* **596**, 583–589 (2021).

9.	Li, Z. *et al.* Structural and dynamic basis of DNA capture and translocation by mitochondrial Twinkle helicase. *Nucleic Acids Res.* **50**, 11965–11978 (2022).
